# Designing symmetrical multi-component proteins using a hybrid generative AI approach

**DOI:** 10.1101/2024.06.13.598662

**Authors:** Delphine Dessaux, Samuel Buchet, Lucie Barthe, Marianne Defresne, Gianluca Cioci, Simon de Givry, Luis F. Garcia-Alles, Thomas Schiex, Sophie Barbe

## Abstract

Proteins, the fundamental building blocks of biological function, orchestrate complex cellular processes by assembling into intricate structures through meticulous interactions. The design of specific protein-protein interfaces to create customized protein assemblies holds immense potential for various biotechnological applications. To address the current limitations in designing heteromeric interactions for multi-component assemblies, we developed a hybrid generative AI design approach. This method combines deep learning and automated reasoning, explicitly considering both positive and negative interaction states to favor heteromeric desired over undesired interactions. The approach leverages Effie, a deep-learned pairwise decomposable scoring function, and an advanced reasoning tool extended for multicriteria optimization of this function. Here, we tested the ability of this hybrid AI method to redesign homomeric interfaces of bacterial microcompartment components (BMC-H) into heteromeric assemblies. We benchmarked its performance against ProteinMPNN, a sequence design autoregressive model. Our in silico assessment, complemented by experimental validation, highlights the outperformance of the hybrid AI generative design approach, and its potential to unlock the engineering of complex multi-component self-assembling protein entities.

## Introduction

Protein-protein interactions (PPIs) play a fundamental role in the functioning of biological systems, governing a wide range of cellular processes.^1,2^ Proteins possess the remarkable ability to self-assemble in highly ordered and functional architectures^3,4^ essential for various activities such as multivalent binding, ultra-sensitive regulation, or compartmentalization. This ability makes the reprogramming of self-assemblies a focal point of extensive research,^5–11^ given their transformative potential in biotechnological applications, including the spatial organization, encapsulation, and transport of active biological entities like enzymes and small molecules.

In nature, protein assemblies often exhibit a significant degree of symmetry,^12^ with the majority being homomeric and consisting of several copies of a single protein monomer sequence. A much smaller fraction are heteromeric assemblies, (i.e., multi-component systems with two or more different protein monomer sequences) which can perform diverse functions that are unachievable with homomeric assemblies alone. The capability to redesign homomeric assemblies into heteromeric assemblies opens new avenues for the development of more sophisticated protein-based entities and novel applications in synthetic biology, nanotechnology and biomedicine.

In protein assemblies, the interactions between the monomers (or subunits) critically determine the collective behavior of each assembly. Achieving high interaction specificity is therefore crucial for the proper functioning and architecture of protein assemblies. This involves ensuring that each subunit of the assembly specifically interacts with its intended target subunit (referred to as the “positive state”) while disfavoring interactions with nontarget subunits (referred to as “negative states”). The challenge of designing such specificity becomes particularly acute in assemblies that should have distinct sequences but should share a similar fold. This arises when the objective is to redesign homomeric assemblies into heteromeric assemblies that must maintain the same symmetric architecture but incorporate distinct sequences (such as A and B). These sequences must retain their fold and form heteromeric interactions (AB, positive state) while avoiding homomeric interactions (AA or BB, negative states, see Figure 1). Consequently, the explicit consideration of these undesirable interactions (negative states) through a multi-state design approach becomes unavoidable.^13^

**Figure 1:**
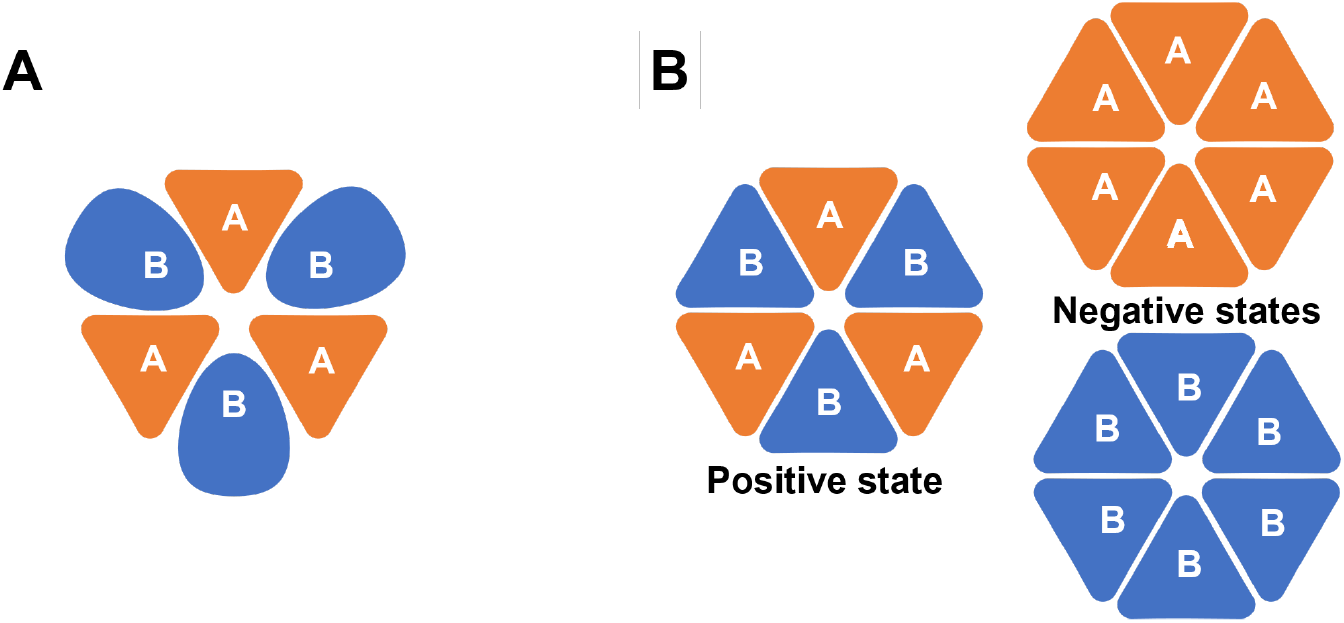
**A)** In a two-component assembly where each component uses a different fold, the identification of two different sequences for the two components is naturally guaranteed by the requirement of having different folds. This guarantee is lost when all components share the same fold. **B)** Designing two orthogonal components with the same fold requires explicitly representing the two undesirable homomeric negative states on the right.

In physics-based approaches to multi-component protein design, such as Rosetta,^8,14–16^ specificity is implicitly introduced by favoring complex interfaces, where affinity requires a precise atomic organization in hydrogen-bond networks and salt bridges. Explicitly accounting for negative states is challenging, not only because exhaustively enumerating negative states is difficult, but also because negative design pits the designer against the laws of physics: the sequence designer tries to destabilize negative states, but physics favors stable conformations, especially when reorienting side chains. This makes negative design computationally excruciating.^17^

In the last few years, deep learning (DL) has revolutionized protein design.^18–25^ DL structure-to-sequence models such as GVP-GNN, ^26^ and ProteinMPNN^22,27^ have shown an impressive capacity to design stable and expressible proteins. These models can be used for design, by sampling the learned conditional probability *P* (*sequence* | *structure*), or for scoring sequences by computing a −*log*(*P* (*·*|*·*)) score, that matches large probabilities to low scores. To differentiate these two usages, we use ProteinMPNN^*D*^ when designing and ProteinMPNN^*S*^ when scoring.

The ProteinMPNN model ^22^ has been used to design sequences for two-component assemblies using components, with different folds.^8^ ProteinMPNN is autoregressive. For design, it successively samples each amino acid from a learned conditional probability distribution represented in logspace, as logits. For multi-state design, a dedicated script uses a linear combination of the logits produced on the various considered states to sample each amino acid.^22^ For negative design, the resulting linear combination captures conflicting objectives, trying to minimize and maximize scores defined over the same fold, for each amino acid sampled. The resulting distribution may be very different from the learned distribution, which may lead to suboptimal sampling.

To address this issue, we developed a hybrid generative AI approach combining Effie, a recent deep-learned pairwise decomposable score function for sequence design,^28^ with an automated reasoning design tool capable of optimizing this function. Because of its pairwise decomposable nature, Effie is similar to pairwise decomposable physics-based score functions, as available in Rosetta. However, Effie is coarse-grained: it has implicitly learned how physics organizes side chains. Negative design with Effie avoids playing against physics and therefore avoids the associated excruciating computational complexity.^17^ This reduces the complexity of negative multi-state design by one level in Stockmeyer’s Polynomial Hierarchy. ^29^ Effie is also far more accurate than Rosetta’s full-atom scoring function in reconstructing the sequence of natural proteins.^28^ It also improves over the related TERMinator score function,^30^ with a more efficient architecture and an enhanced loss function.^28^ The resulting hybrid architecture combines the accuracy of deep learning with the ability of automated reasoning engines to satisfy extra constraints capturing design requirements, without the limitations of black-box autoregressive models. For this work, we extended the automated reasoning engine toulbar2^31,32^ used in Custozyme+, our design software, to handle multiple criteria (states). Effie can be optimized to design sequences or can directly score sequences. We use Effie^*D*^ and Effie^*S*^ to differentiate these two usages.

Inspired by the intricate PPIs within Bacterial Microcompartments (BMCs), we applied these approaches to redesign homomeric BMC components (Figure 2). BMCs stand out as prime illustrations of nature’s adeptness at structuring elaborate biomolecular architectures via meticulous PPIs. These proteinaceous entities, found in the cytoplasm of many bacteria, comprise an outer shell enclosing enzymes involved in specific metabolic pathways (e.g., the degradation of 1,2-propanediol or ethanolamine, in the PDU or the EUT respectively, or the fixation of CO_2_ by carboxysomes) and scaffolding components.^33–42^ Notably, BMC shells contribute to metabolic pathway optimization and limit the diffusion of volatile or toxic reaction intermediates.^43^ Beyond their function as natural nanoreactors, appealing for synthetic biology,^44–46^ the extraordinary self-assembly properties of BMC shell components justify a strong interest for varied bio(nano)technological aims.^33,41,42,47,48,48–54^

**Figure 2:**
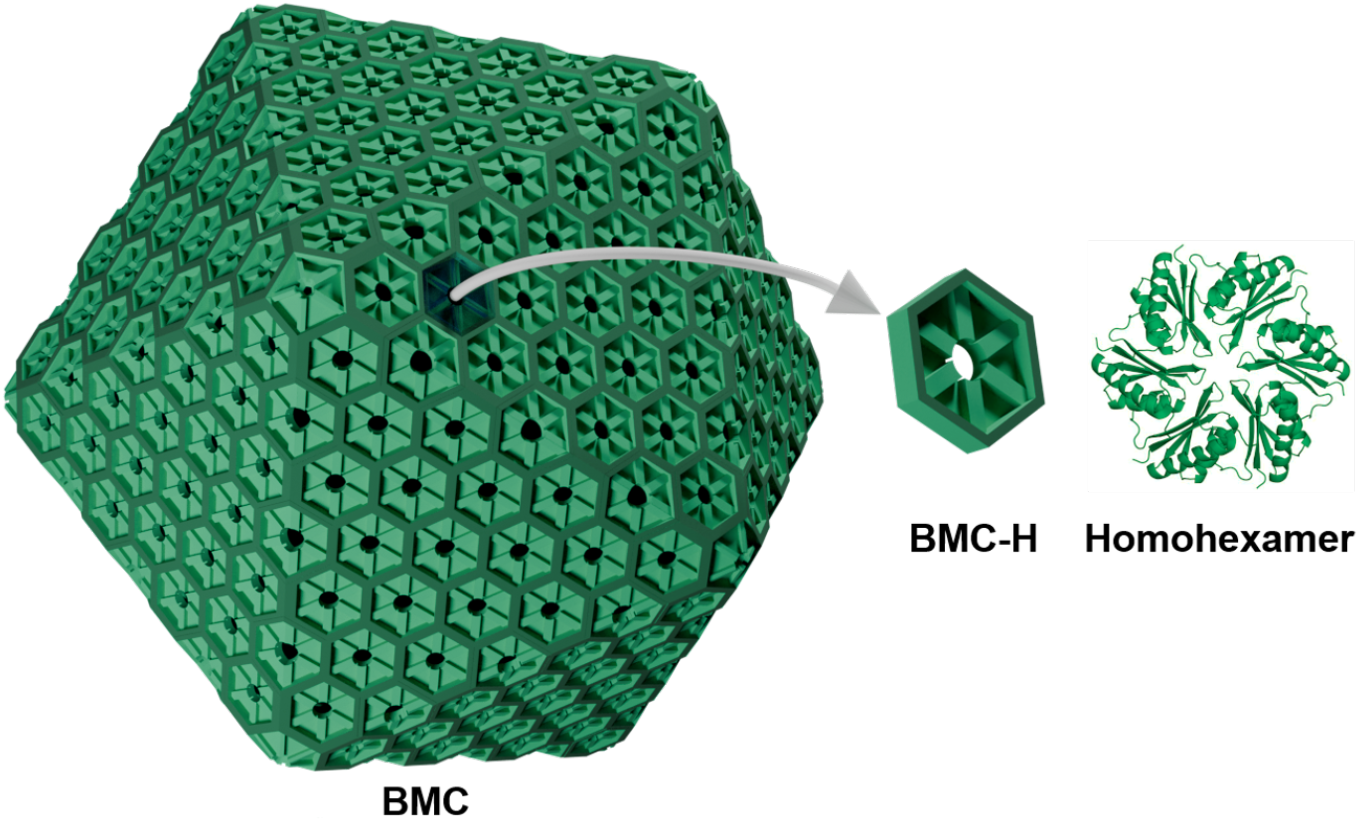
BMC-H homohexamer of bacterial microcompartment (BMC) shell.

The most abundant BMC-shell component is a hexameric assembly known as BMCH. When expressed alone, BMC-H can self-assemble into a variety of higher-order objects including nanotubes, spheroids, Swiss-rolls, or flat sheets.^47,55–58^ To better exploit BMC shell components for bio(nano)technological applications, their precise organization would need to be controlled, and more specifically, the position of each subunit should be precisely adjustable. For this, designing a modular set of proteins that self-assemble in a controlled ordered symmetric assembly would be needed. Such control over the position of the different BMC-H monomers within the hexamer could eventually serve to tune the spatial organization of enzymatic modules that could be covalently linked to these monomers with precision. One first step in this direction consists in specializing the monomer of BMC-H into two sequences 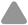 and 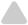 that should assemble into the positive hexameric state 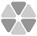, but avoid the negative homomeric states 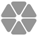 and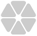 (see Figure S1).

In this study, BMC-H interfaces were thus redesigned using either Effie^*D*^, implemented in Custozyme+, or ProteinMPNN^*D*^ to form heteromeric interactions. The design results obtained using both methods were assessed in silico by cross-evaluating every designed sequence pair AB using both Effie^*S*^ and ProteinMPNN^*S*^. Twenty-four AB sequence pairs generated either with Effie^*D*^ or ProteinMPNN^*D*^, were experimentally tested using a tripartite GFP construct^59,60^ to check for the heteromeric interaction between both monomers. Fluorescence signals and co-purification of the monomers along with the GFP partner were used to select the best candidates for further characterization. Eventually, heterohexamer formation was verified for the best-designed sequences using chromatography approaches.

## Results and discussion

The first result of this work is the definition and implementation of a hybrid AI method for multistate protein design, including possible negative states, which combines Effie, a deep-learned pairwise decomposable score function predictor^28^ with a multicriteria extension of `toulbar2`,^31,32,61^ a discrete optimization prover based on automated reasoning, with a nice record in provable computational protein design.^62^ While this hybrid approach may be computationally more demanding than sampling from an autoregressive model, it allows for the simultaneous optimization of multiple states. This hybrid approach directly tackles the multi-state design problems. Instead, the heuristic linear combination of autoregressive deep learning models proposed in ProteinMPNN^22^ struggles to sample sequences optimizing the conflicting objectives of negative multi-state design.

### Comparison between Effie and ProteinMPNN results

Pairs of sequences 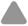 and 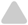 were designed so that all these sequences fold onto the structure of a monomer of the RMM (*Rhodococcus* and *Mycobacterium* microcompartment) BMC-H, and self-assemble as a heteromeric assembly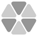, which defines the positive state (Figure 1B). To prevent the sequences from assembling into the homomers 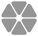 or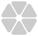, these two states were considered negative states. In the first approach, the design was performed by Effie^*D*^. In the second approach, another set of sequences 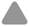 and 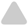 was designed applying the dedicated multistate design helper script provided with ProteinMPNN.^22^ Both approaches are described in detail in the Materials and Methods section.

In each approach, a symmetrical homohexamer conformation of RMM was used as a template for designing pairs of sequences. The design was focused on the residues located at the interface between the monomers in the assembly. Two designable regions were considered: a small region consisting of 29 residues directly at the interface, and a larger region of 39 residues, including nearby residues (see Figure S2).

To evaluate and compare the design results, all predicted sequences were scored on the positive and negative states, with Effie^*S*^ and ProteinMPNN^*S*^, defining *pre-min* scores. Following minimization with Rosetta `beta nov16` function,^63^ they were re-evaluated with both score functions, defining *post-min* scores. The differences between the scores obtained on the positive and each negative state 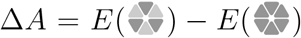 and 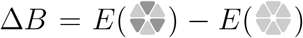 were computed for each design. They indicate the effectiveness of the design in satisfying the objective of favoring positive over negative states. The difference between the score of the designed heterohexamer 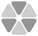, and the score of the wild-type (WT) homomeric RMM template was also computed with both score functions, denoted as 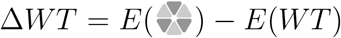, to assess how far the designs deviate from the wild-type.

To account for possible discrepancies in the score units of Effie^*S*^ and ProteinMPNN^*S*^, score differences were normalized by the standard deviations over all Δ*A* and Δ*B* values for Effie^*S*^ or ProteinMPNN^*S*^ 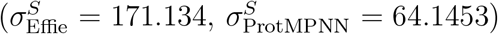. For a design scored with Effie^*S*^, this defines 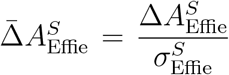. Normalized score-differences 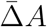 and 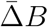 obtained for designs for the small design region are represented in Figure 3.

**Figure 3:**
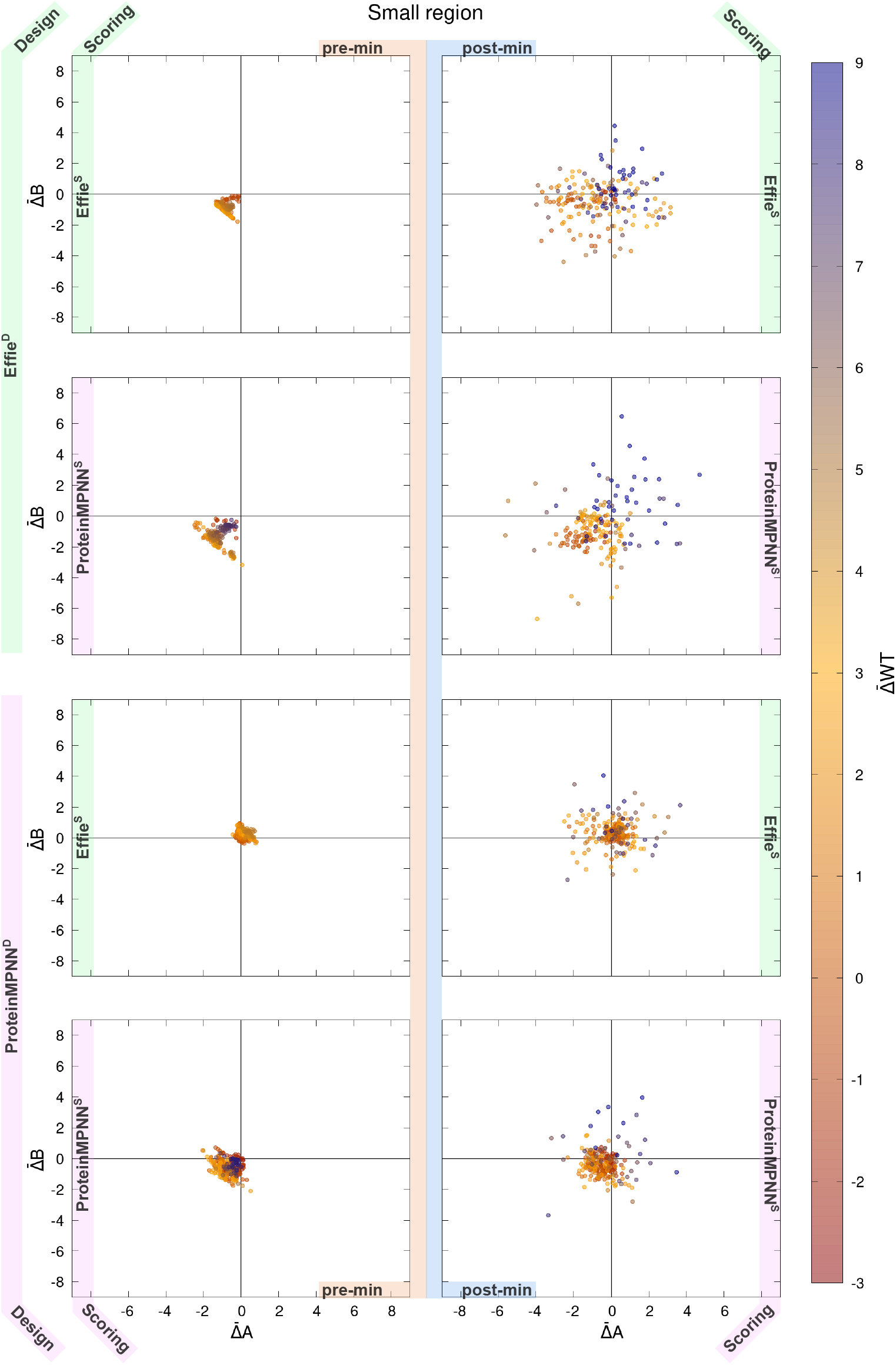
Comparison of the score differences between positive and negative hexameric states for mutants generated by Effie^*D*^ or ProteinMPNN^*D*^ on the small design region. The left column represents pre-min scores and the right column represents post-min scores. Every XY dot-plot represents the 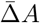 and 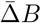 on *X* and *Y* axis. Each dot represents a design with its color indicating the Δ*WT* score: blue colors indicate a likely weakly stable 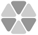, compared to the WT. The two top rows show Effie^*D*^ designs, while the two bottom rows show ProteinMPNN^*D*^ designs. Scoring is alternatively performed using Effie^*S*^ (top and 3rd rows) and ProteinMPNN^*S*^. Focusing just on ProteinMPNN^*S*^ scores, Effie^*D*^ designs can simultaneously have better 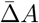 and 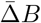 (be in the bottom left corner) while having a good score compared to the WT. The best design of ProteinMPNN^*D*^ in terms of 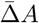 and 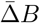 have a poor score compared to the WT.

In our design problem, scores for the positive state 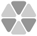 should be lower than scores for the negative states 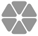 and 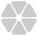: the lower 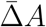 and 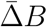, the better. Similarly, to ensure the stability of the heterohexamer 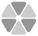, its score should not exceed the template score by too much: the value of 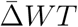 should be close to 0 or negative.

As shown in Figure 3, for the small design region, designs 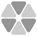 generated with Effie^*D*^ exhibit overall better score differences 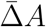 and 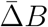 between positive and negative states than those of ProteinMPNN^*D*^, regardless of the function used for scoring. This means that ProteinMPNN^*S*^ prefers the sequences designed by Effie^*D*^ over those designed by ProteinMPNN^*D*^, confirming that autoregressive models have difficulties sampling high-probability regions in constrained settings. This trend is particularly pronounced before minimization.

To numerically assess this tendency, we define for each design *i* the vector 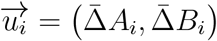.The further this vector extends in the direction [−1, −1], the better. For each design method, we hence computed the average scalar product on pre and post-minimization scores:

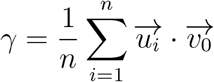

with 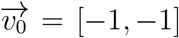. Table 1 provides *γ* values for all scoring evaluations with a higher *γ* indicating better design results.

**Table 1:** Evaluation of Effie^*D*^ or ProteinMNN^*D*^ designs for each designable region. (*n*) is the number of designs generated, (*γ*) indicates how much the positive state is improved over the negative states (see main text), (yield) gives the percentage of in silico successful designs, where the positive state gets a better score than either negative states.

| Scoring function | | → | Effie<sup>S</sup> | | | | PMPNN<sup>S</sup> | | | |
| --- | --- | --- | --- | --- | --- | --- | --- | --- | --- | --- |
| Design function | Design space | $n$ | pre-min | | post-min | | pre-min | | post-min | |
| | | | $\gamma$ | yield | $\gamma$ | yield | $\gamma$ | yield | $\gamma$ | yield |
| Effie<sup>D</sup> | small | 189 | 1.75 | 100 % | 0.92 | 40.7 % | 2.58 | 99.5 % | 1.37 | 58.2 % |
|  | large | 188 | 2.52 | 100 % | 0.92 | 48.4 % | 3.16 | 99.5 % | 1.82 | 70.2 % |
| PMPNN<sup>D</sup> | small | 236 | -0.40 | 3.0 % | -0.53 | 10.2 % | 1.23 | 82.6 % | 0.78 | 55.1 % |
|  | large | 259 | -0.68 | 1.9 % | -0.50 | 9.3 % | 1.65 | 88.0 % | 0.93 | 57.5 % |

Before minimization (left column of Figure 3 and S3), the evaluation of Effie^*D*^ designs by ProteinMPNN^*S*^ 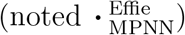 gives a 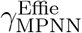 of 2.58 and 3.16 on the small and large designable regions. The evaluation of ProteinMPNN^*D*^ designs 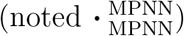 on the same regions, gives lower 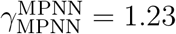 and 1.65, respectively.

Using Effie^*S*^ for evaluation instead, ProteinMPNN^*D*^ designs are considered even worse with 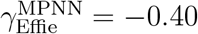 for the small region. In fact, for either designable region, ProteinMPNN^*D*^ designs give a negative 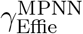, implying that Effie^*S*^ evaluates most of these sequences as not meeting the design objectives. Under the specific design constraints that negative multistate induces, autoregressive sampling clearly struggles to recover sequences satisfying the negative multi-state design objectives.

After minimization, scores vary significantly, resulting in more scattered values for 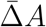 and 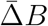 in Figures 3 and S3 (right column). However, in all cases *γ*_post min_ *< γ*_pre min_. This is consistent with the distortions of the hexamer conformations obtained for designs with strong negative weights *λ*^−^, resulting in high scores for states 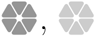, and also 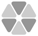 and large differences 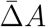 and 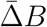 (see Figures S4 and S5). For ProteinMPNN^*D*^ designs, Effie^*S*^ 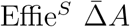 and 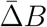 still tend to be positive post-minimization, with negative 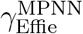 values, as shown in Table 1, with only a few designs displaying negative values for both 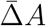 and 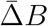 (less than ∼ 10 % of designs for either region).

ProteinMPNN^*S*^ scoring shows more designs with negative 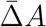 and 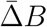, but 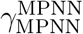 remains lower than 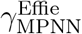 of Effie^*D*^ sequences. A possible explanation of the lesser degradation of ProteinMPNN^*D*^ designs scores after minimization lies in the different amounts of noise used during training between ProteinMPNN (0.2 Å std-dev) and Effie (0.14 Å std-dev), suggesting that using larger amounts of noise for Effie training could be useful.

All post-minimization *γ* values increase with the size of the design region which shows that both methods are more successful at negative multistate design on larger design regions.

While this is expected for the optimization-based Effie^*D*^, this shows that autoregressive sampling, despite its limitations here, is still able to exploit additional design space.

Also, in all cases, *γ*_Effie_ *< γ*_MPNN_, which highlights the fact that ProteinMPNN tends to be less stringent than Effie in its design evaluation. On average, Effie^*D*^ designs contain 58*±*10 mutations whereas ProteinMPNN^*D*^ designs have 49*±*12 mutations. Hence, it appears that the higher number of mutations integrated into the sequence via Effie^*D*^ provides better results for our design problem.

Another advantage of Effie is that it predicts a pairwise decomposable scoring function instead of just a black-box auto-regressive score. It is therefore possible to separate the score of a multimeric structure into intra and inter-chain scores. Effie thus provides valuable insights into the effect of mutations on the interaction between chains. Figures S6 and S7 display the differences between inter-chains scores of the heteromer 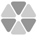 and the homomers 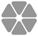or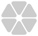, i.e., 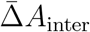 and 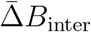, and Table S1 summarizes the scalar product *γ*_inter_ (calculated using 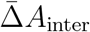 and 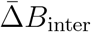 normalized by the same *σ*_Effie_ used to normalize values calculated on total Effie^*S*^ scores). The score differences between positive and negative states are mainly due to differences in inter-chain scores, as *γ*_inter_ ≈ *γ*_Effie_. The ΔWT inter-chain scores show that ProteinMPNN^*D*^ heteromeric designs scores are closer to the WT score than Effie^*D*^ designs (see Figures S6 and S7). However, the 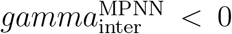*<* 0 show that ProteinMPNN^*D*^ designs rarely favor the heteromeric state 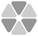 over the homomeric state 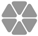 and 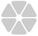 (see Table S1), failing to optimize the objective of multistate negative design.

### Experimental assessment

For a more extensive comparison of Effie^*D*^ and ProteinMPNN^*D*^, pairs of sequences generated with both approaches were selected for experimental testing. This selection was mainly based on the score differences, Δ*A* and Δ*B*, calculated with the respective scoring functions used for design. In addition, a diversity criterion was also applied, with a minimum of 10 mutations between sequence pairs required for selection. Therefore, 24 sequence pairs were retained for experimental testing (see List S1 for details on amino acid sequences and numbers of mutations), 14 from Effie^*D*^ (Duo1 to 14) and 10 from ProteinMPNN^*D*^ (Duo15 to 24). The score differences, Δ*A*, Δ*B* and Δ*WT*, calculated using either Effie^*S*^ or ProteinMPNN^*S*^ are detailed for each of the 24 selected designs in Tables S2 and S3. Table S4 reports the inter-chain Effie^*S*^ score differences.

Interactions between both members from these 24 selected sequence pairs, were screened using the tripartite green fluorescent protein (tGFP) technology.^59^ This technology was recently adapted in our laboratory for the study of BMC shell subunits oligomerization^60^ and is based on the splitting of the 11-stranded GFP reporter into three non-fluorescent portions: the large GFP1-9 portion, plus the two terminal *β*-strands GFP10 and GFP11 that are expressed in fusion with the two proteins to be assayed. Reconstitution of the complete GFP reporter, with concomitant fluorescence emission, is promoted by the association of the two protein partners, whereas the entropic penalty induced by the encounter of three partners hampers fortuitous GFP reconstitution (see Figure S8A).

The DNA sequences that code for each pair of GFP10/11-labeled proteins were engineered as bicistron in a plasmid also permitting independent transcription of an oligo-histidine-tagged GFP1-9. After transformation of BL21(DE3) *E. coli* cells, fluorescence emission was monitored over the course of cultures, with an induction of protein expression upon inoculation of the culture. After fitting data to a sigmoidal function, maximal fluorescence values (*F*_*max*_) from several replicates were averaged to generate data presented in Figure 4A (see Materials and Methods for further details).

**Figure 4:**
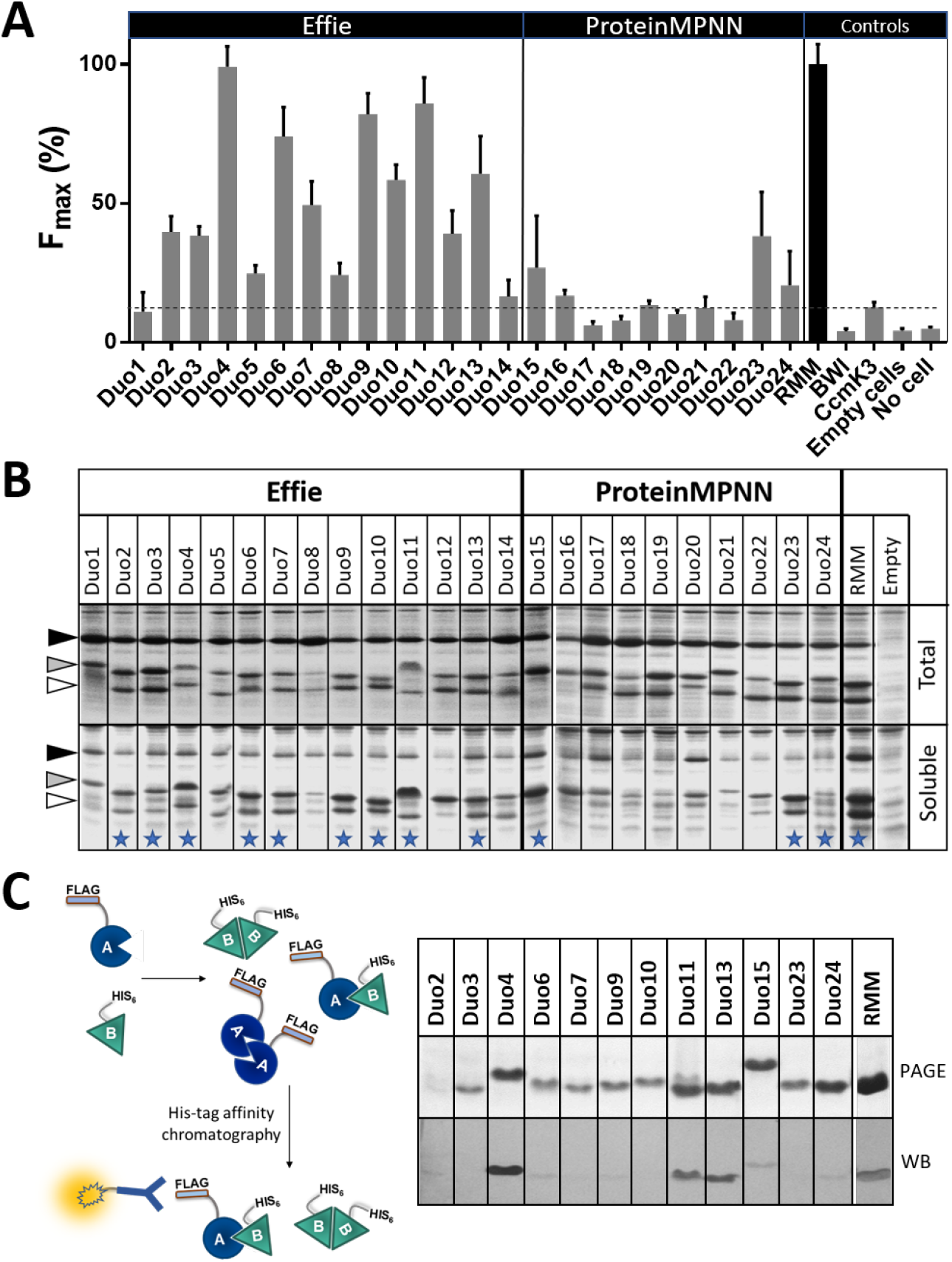
Experimental behavior of the Duo designs.**A)** Screening of the associations between designed pairs of monomers A and B using the tripartite GFP technology. Maximal fluorescence (*F*_*max*_) values were estimated from fits to a sigmoidal function of the GFP fluorescence data monitored during growth of *E. coli* bacteria overexpressing each Duo (see Materials and Methods for details). Data are given as a percentage of the WT RMM value (i.e., two identical monomers connected to GFP10 and GFP11 peptides). The dashed line highlights maximal signals emitted by negative controls: non-associating pairs (BWI), aggregating proteins (CcmK3 from *Syn. Sp. PCC 6803*) or cell auto-fluorescence (Empty: cells transformed with a pET26b empty vector). **B)** Expression and solubility of screened designs. Cells overexpressing A-GFP10 and B-GFP11 monomers, plus the GFP1-9, were recovered 4 h post-induction (see M&M). Total cellular contents (top) and proteins remaining soluble after lysis and centrifugation (bottom) were analyzed after thermal denaturation on SDS-polyacrylamide gels (see also Figure S8). The white, grey, and black arrows indicate the approximate positions of B-GFP11, A-GFP10 and GFP1-9, respectively. **C)** Selected cases (indicated with a star on B were further investigated using the approach depicted on the left: monomer A is connected to a Flag peptide, whereas monomer B is fused to a His_6_-purification tag. After overexpression and cellular lysis, soluble proteins carrying the purification tag are retained inside cobalt-based chelating resins. After washing, bound material is eluted from the resin and analyzed through Coomassie blue-stained polyacrylamide gels (right side, Top). Detection of co-purified A-FLAG partners is performed in parallel by Western Blot, using Flag-specific antibodies (Bottom). Only the portion of the gel corresponding to the region of migration of the Duo monomers is shown for clarity.

Globally, expression of Duo1 to Duo14, designed with Effie^*D*^, generated a stronger fluorescence than Duo15 to Duo24, deriving from ProteinMPNN^*D*^. All Effie^*D*^ pairs, omitting Duo1 and to a lesser extent Duo14, reached fluorescence levels clearly above those of the negative controls CcmK3 and BWI (threshold highlighted by a dashed line on Figure 4A). Moreover, the *F*_*max*_ of 6 out of the 14 Effie^*D*^ designs was above 50 % of values recorded for the WT RMM, our positive control reference. On the contrary, signals for 7 ProteinMPNN^*D*^ pairs were below, or comparable to the negative control threshold, and none of the 10 designs attained a *F*_*max*_ comparable to the one of the WT RMM.

Analysis of protein contents by SDS-PAGE provided additional proof for the superiority of Effie^*D*^ designs (see Figure 4B). Although the expression levels of A-GFP10/B-GFP11 monomers and GFP1-9 were found to be adequate for the majority of the pairs, regardless of the design method, band intensities in soluble fractions appeared to decrease more often and more severely for ProteinMPNN^*D*^ pairs. These soluble fractions were prepared by centrifugation after cellular lysis, which induces the sedimentation of aggregated material, membranes, and other cellular debris. Aggregation of a fraction of the expressed BMC-H proteins therefore likely explains this decrease in band intensities mainly observed for ProteinMPNN^*D*^ designs. Of note, purification of a selection of cases using cobalt-based affinity chromatography proved that the two GFP10 and GFP11-tagged partners co-eluted in association to His_6_-tagged GFP1-9 (Figure S8B).

Nine Effie^*D*^ and three ProteinMPNN^*D*^ designs were further investigated using a different experimental approach based on the attachment of a Flag peptide and the His_6_ tag on monomers A and B from each pair, respectively (Figure 4C). The cross-association of both monomers from the Duo4, Duo11, and Duo13 was in that manner proven by the antibody-based detection of the Flag-carrying monomer co-purifying with the His_6_-tagged monomer.

Moreover, the western blot band intensities of these three designs are comparable or even higher than the ones detected for the positive control, consisting of a combination of Flag- and His_6_-tagged WT RMM monomers (Figure 4C).

Duo4 yielding the most intense western blot band on Figure 4C, was characterized in more detail. Namely, after purification, its oligomeric state was investigated by size-exclusion chromatography (SEC), which revealed the existence of two preponderant species: one eluting at volumes expected for a hexamer and the second migrating as a dimer (see Figure S9A). Some aggregated material was also detected. These two major peaks were collected separately from each other and, after reconcentration, they were re-injected in the SEC column under the same conditions, which preserved their respective elution volumes (i.e., oligomeric state), as observed in Figure S9B. More importantly, each oligomeric state remained stable under the conditions of our experimental setup, ruling out equilibration between the two oligomeric species in solution. Therefore, we hypothesize that the formation of dimers (or aggregated species) might be the consequence of phenomena not directly related to the protein structural design problem, such as the difficulties of precisely tuning the expression levels and rates of each monomer inside the cells, ^64^ which could be regulated by using optimized genetic organizations.

Overall, these results show that Effie provides a more appropriate description of multidimensional landscapes assessed in negative design problems of such new associations of pairs of mutant BMC-H proteins than ProteinMPNN.

## Conclusion

For decades, computational protein designers have relied on physics-based approaches such as Rosetta,^14^ combining rigid backbones assumptions, flexible and mutable side-chains represented as rotamers, and a pairwise decomposable energy function that was minimized using either stochastic Monte-Carlo approaches or provable optimization methods such as DEE/*A*^*\** 62,65,66^ or automated reasoning approaches ^17,61,67–70^ as embodied in the Cost Function Network solver toulbar2. While computationally expensive, this approach allows for various a posteriori constraints that could, for example, enforce the existence of hydrogenbond networks^71^ or avoid specific amino acid types or combinations of them in the designed sequence.

The advent of deep learning-based approaches has revolutionized computational protein design. For sequence design, autoregressive models can give direct access to high-probability samples of sequences *s* given a state *B P* (**s**|*B*) by iteratively sampling the learned *P* (**s**_*i*_|**s**_*<i*_, *B*) at low temperature, one amino acid after the other. Autoregressive models however, rely on the exact chain-rule identity *P* (**s**|*B*) = Π_*i*_ *P* (**s**_*i*_|**s**_*<i*_, *B*) which assumes that future amino acids will be sampled from *P* (**s**_*i*_|**s**_*<i*_, *B*). This is no longer true if sampling is done from a constrained or biased distribution instead of the learned *P* (**s**_*i*_|**s**_*<i*_, *B*). Even if we assume that the trained neural network has perfectly learned *P* (**s**_*i*_|**s**_*<i*_, *B*), the guarantee that top-probability sequences will be sampled at low temperature is lost.

As an alternative to autoregressive models, deep learning can be used to predict a coarsegrained decomposable score function *E*(**s**|*B*) ∝ *P* (**s**|*B*), as does Effie.^28^ While computationally more expensive to sample than autoregressive models, decomposable pairwise models (aka Potts or Markov Random Field models^30,72–74^) can be arbitrarily constrained or biased and then optimized to produce highest-probability sequences. As this paper shows, pairwise decomposable score-based designs are considered better than those produced by a state-of-the-art autoregressive model, by the auto-regressive model itself, thereby proving its own failure to sample high probability regions.

In our attempts at redesigning the monomer of a BMC-H into two different sequences, 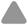 and 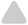 assembling into 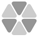, this intrinsic weakness extends to experimental results, as most autoregressive-predicted sequences tested did not reveal structurally viable, as demonstrated using in vivo (tripartite GFP) and in vitro (western blots) screens. Instead, our hybrid approach, which combines Effie, ^28^ a deep-learned decomposable score function, with toulbar2,^31,75^ an automated reasoning solver, was able to produce pairs of sequences that not only caused strong fluorescence in vivo or western blot band intensities in vitro, but most remarkably were also soluble and formed heterohexamers. These results pave the way for building a catalog of six monomer sequences that would self-assemble into a uniquely defined BMC-H hexamer, itself able to self-assemble into arrays, nanotubes, or other higher-order assemblies.

## Materials and Methods

We use bold letters to denote sequences of objects (e.g., **s** for a sequence of amino acids). Each sequence’s element is denoted by a plain letter such as *a* ∈ **s**. The element at position *i* in a sequence **s** is denoted as **s**_*i*_. Assuming an implicit order over residues, **s**_*<i*_ and **s**_*>i*_ will denote the sequences of amino acids respectively before and after position *i*.

### Multistate design with Effie and bi-objective optimization

We assume that we have a set of positive and negative rigid backbone states **B** = **B**^+^ ∪ **B**^−^, all with the same number *ℓ* of residues. At each position 1 ≤ *i* ≤ *ℓ*, we have a set *S*_*i*_ of possible amino acids.

Given a state *B*, and a sequence **s**, Effie has been trained using an improved variant of the pseudo-likelihood loss function^28^ to predict a score function *E*(**s**|*B*) such that the probability *pP* **s**|*B*) of the sequence **s** given the state *B* satisfies:

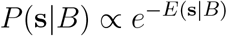

Given a state *B*, the neural architecture of Effie pr edicts th e sc ore fu nction *E*(**s**|*B*) as a pairwise decomposable score function, i.e., a sum of terms that each involve two residues:

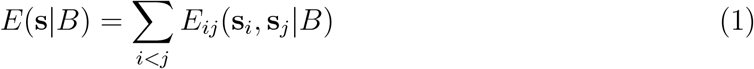

Each pairwise term is predicted as a 20 *×* 20 tensor (matrix). No constant or unary term needs to be predicted, as these can be implicitly represented in pairwise terms.^74^

When designing multi-component symmetrical proteins, the identity of a given component’s *i*^*th*^ residue will be the same in all its occurrences in each sub-unit of the symmetrical assembly. In (1). Every variable that does not represent the first occurrence of a component can therefore be replaced by the corresponding variable in the component’s first occurrence. This drastically reduces the number of variables in the above sum. For a C6 monomeric hexamer, such as 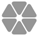 or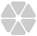, this reduces the number of variables by 6. For a two-component C3 hexamer, such as 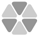, the number of variables is reduced by 3. These reductions are applied systematically in the rest of the paper.

The multistate sequence design (MSD) problem is to find a sequence **s** which minimizes:^17^

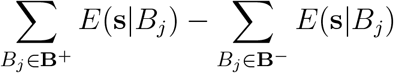

Indeed, minimizing this difference means minimizing the score of the designed sequence **s** on positive states and maximizing it on negative states, thus favoring positive against negative states.

Given that the score function *E* predicted by Effie is pairwise decomposable, this optimization problem is known to be NP-complete, being mathematically equivalent to the usual single-state protein design problem, ^76^ with each term being the difference of terms between positive and negative states. It is however considerably simpler than the full-atom physics-based formulation which was proven to be at the second level of Stockmeyer’s Polynomial Hierarchy (i.e., NP^NP^-complete or 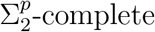).^17,29^ This objective can be optimized using any available discrete pairwise decomposable energy minimization tool. We use the Cost Function Network solver toulbar2^31^ for its efficiency^62^ and the variety of algorithms it implements, including both systematic provable optimization algorithms ^77^ and stochastic optimization algorithms.^78^

To more precisely tune the contribution of positive versus negative states, we assume that weights *λ*^+^ and *λ*^−^ are given (equal to respectively 1 and −1 in the above formulation), and we are interested in minimizing the weighted score:

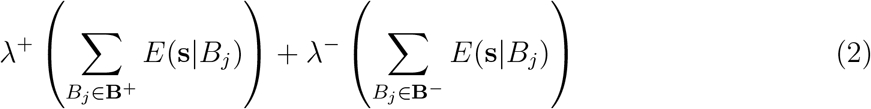

This corresponds to the scalarized formulation of a bi-objective optimization problem (see Supplementary Information, Section 1.1) where the first objective, to be minimized, is 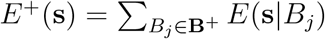 and the second objective, to be maximized, is 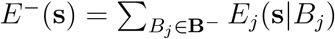, defining a pair of scores for every sequence **s** in the design space. This pair is hereafter denoted as the score-pair (*E*^+^(**s**), *E*^−^(**s**)).

In this context, a given sequence **s** is said to dominate another sequence **t** (i.e., “**s** is better than **t**”) if **s** has a strictly better score for one objective and an equal-or-better score for the other one, i.e., either *E*^+^(**s**) *< E*^+^(**t**) and *E*^−^(**s**) ≥ *E*^−^(**t**), or *E*^+^(**s**) *≤ E*^+^(**t**) and *E*^−^(**s**) *> E*^−^(**t**).

Two sequences that do not dominate each other are considered incomparable. The goal of multi-objective optimization is to find a set of incomparable sequences that are not dominated by any other sequences. By varying the weights (*λ*^+^, *λ*^−^), it is possible to obtain a set of non-dominated sequences whose score-pairs correspond to the convex hull of the set of all non-dominated score-pairs, the so-called Pareto front.^79^ When *λ*^+^ *>* 0 and *λ*^−^ *<* 0, the sequences minimizing (2) are guaranteed to be non-dominated solutions, as shown in Section 1.1 in the Supplementary Information.

### Approximation of Pareto frontiers

While only NP-complete, exactly optimizing the scalarized score (2) defines a still challenging problem, even if one relies on the efficient provable algorithms implemented in the Cost Function Network solver toulbar2.^17^ In the mono-objective case, two types of approaches can be used when proving optimality is unfeasible in reasonable time:

1. One can forcibly stop systematic *provable* optimization tools before they finish their proof. In this case, a current best (incumbent) solution and a lower bound on the optimal cost are produced. The lower bound and the cost of the incumbent solution together define an optimality gap.
2. Instead, one can rely on *stochastic* optimization methods such as Monte Carlo Simulated Annealing, or Large and Variable Neighborhood Search (VNS^78,80^) algorithms that only produce an incumbent solution. Combined with the previous approach, this can tighten the previously determined optimality gap by decreasing the cost of the incumbent solution.

In the bi-objective case, one can similarly approximate the Pareto front by computing a set of incumbent solutions minimizing the scalarized score (2) and a lower bound curve, providing guarantees that the produced incumbent solutions are not too far from optimality.

To approximate the Pareto front, our approach relies on a variant of the dichotomy algorithm^81^ which we call the approximation dichotomy algorithm. The original dichotomy algorithm repetitively optimizes the objective (2) using different weights to perform an exhaustive enumeration of non-dominated score pairs, starting from extreme weights pairs (1, 0) and (0, 1). Each time a new provably optimal solution is found, an intermediate weight pair is computed, and the search is repeated until all possible non-dominated score pairs obtainable by optimizing (2) are produced.

We modified this method to account for potential sub-optimal solutions returned after solving with a given pair of weights. Several conditions were added to control the progression of the Pareto front approximation and ensure the termination of the procedure, as detailed in Section 1.2 of the Supplementary Information. For given weights (*λ*^+^, *λ*^−^), the single-objective (2) can be optimized using both interrupted exact optimization and stochastic optimization, providing a lower/upper bound pair (*lb, ub*) on the scalarized cost. We then know that:

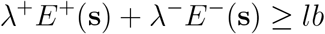

which defines a half-space in which we know that no solution exists. As we repeatedly optimize the scalarized score (2) in the approximation dichotomy, the union of these half-spaces defines a lower bounding region, whose frontier is represented as an orange line in Figure 5. Together with incumbent solutions, they delineate an approximation of the Pareto front in which the designer can choose sequences.

**Figure 5:**
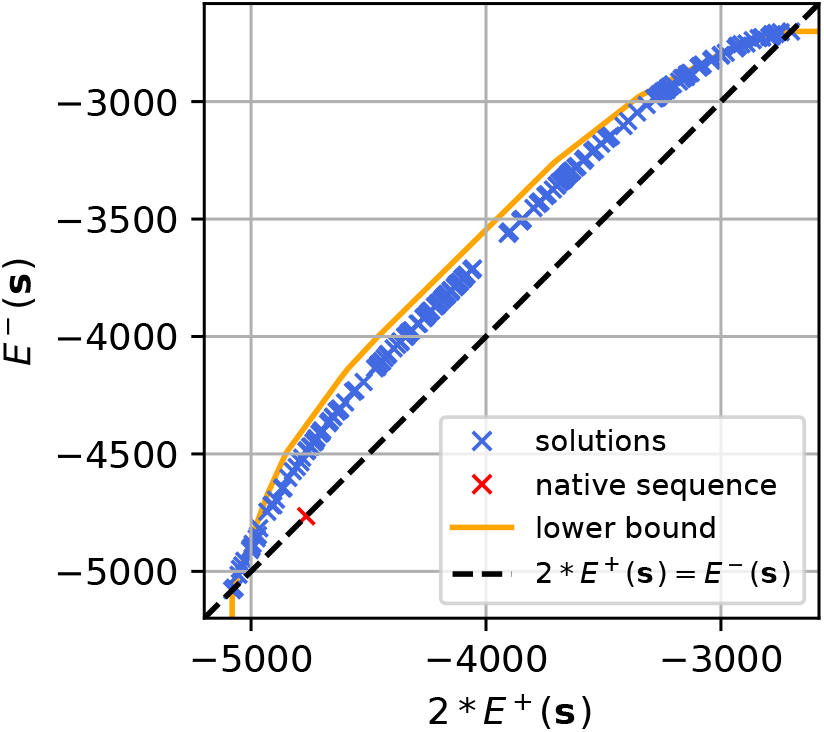
Solutions and lower-bound curve obtained with the approximation dichotomy algorithm. The Pareto front of the problem must be located between the blue points and the yellow curve. The slightly convex curve above the diagonal indicates to which point the scores of the positive state (X axis) and negative state (Y axis) can be pushed away. The native state appears on the diagonal as a red cross. Its position indicates which levels of Effie^*S*^ score should be sufficient to stabilize the considered fold.

The dichotomy algorithm only tries to identify the convex hull of the Pareto front, and the number of solutions needed to define it may be limited. To offer a wider diversity of designs, we generated more incumbent solutions by collecting the solutions produced by the stochastic metaheuristic in toulbar2. Both techniques increase the density of sequences and associated incumbent solutions around the Pareto frontier. To control the number and diversity of the sequences, they are clustered using *MMseqs2*,^82^ adjusting the similarity threshold to the number of desired solutions. A sequence can then be selected in each cluster.

The approximation dichotomy algorithm has been implemented with the Cost Function Network solver toulbar2,^31^ (https://github.com/toulbar2/toulbar2) relying on two algorithms within it: *Hybrid Best First Search (HBFS)* ^77^ was used for exact search, Variable Neighborhood Search (VNS)^78,80^ was used as the metaheuristic.

### Multistate design with ProteinMPNN

ProteinMPNN is an autoregressive model that learned the probability distribution *P* (**s**_*i*_ | **s**_*<i*_, *B*) over amino acids for a residue at position *i* given a state *B* and the identity of previously fixed amino acids *s*_*<i*_. Applied iteratively on a protein with *ℓ* residues, using the chain rule

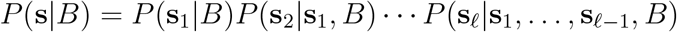

ProteinMPNN can directly sample *P* (**s**|*B*), the conditional probability of sequences **s** given the target state *B*. Using low temperature at each autoregressive step favors the sampling of high-probability sequences.

It is important to note that the marginal conditional probabilities *P* (**s**_*i*_|**s**_*<i*_, *B*) that ProteinMPNN has learned to predict assume that the future amino acids **s**_*>i*_ will also be sampled from the same model. If this is not the case, these predicted marginal conditional probabilities may become arbitrarily wrong and sampling may eventually explore low-probability regions.

For symmetrical design, ProteinMPNN can tie symmetrical residues together. When a tied position is sampled, the shared amino-acid identity is chosen using the sum of the logits of the tied residues.

For multistate design, the authors of ProteinMPNN ^22^ suggest using a linear combination of the logits obtained on different states with positive and negative weights, *λ*^+^ and *λ*^−^, to favor or disfavor chosen states. The GitHub repository for the software (https://github.com/dauparas/ProteinMPNN) contains a helper script `make pos neg tied positions dict.py` for this purpose.

Assuming that a positive state *B*_+_, (weight 1) and a negative state *B*_−_ (weight −1) are used, this means that each amino acid will be sampled from a distribution 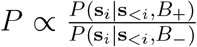 instead of the respective distributions *P* (**s**_*i*_|**s**_*<i*_, *B*_+_) and *P* (**s**_*i*_|**s**_*<i*_, *B*_−_) that ProteinMPNN autoregressive model assumes on either state. This may alter its accuracy in unpredictable ways.

### Implementation of multistate design approaches

#### 3D model construction and definition of designable residues

A 3D symmetrical hexamer conformation was built from the chain A of the homohexamer X-ray structure of the BMC-H shell protein MSM0272 forming the *Rhodococcus* and *Mycobacterium* microcompartment (RMM) from *Mycobacterium smegmatis* MC^2^ 155 (PDB ID=5L38)^83^. After identification of the symmetry axis using AnAnaS,^84^ Rosetta cyclic C6 symmetry definition^85^ was used with a 60° rotation between each asymmetric subunit and a distance of 17.1 Å between the center of mass of the subunits and the center of the cyclic hexamer. The conformation of this hexamer was then minimized using Rosetta `ref2015` Cartesian scoring function^14^ and the same symmetry constraints. The resulting conformation is denoted as the wild-type (WT) template. It was used for the design of heteromers composed of two different subunits, A and B (see Figure S1), and as a reference for design evaluation. A small (29 residues) and a larger (39 residues) designable region were defined at the interface between subunits (see Figure S2).

#### Hybrid AI-based design

Effie was trained using its original architecture^28^ on the multi-chain protein dataset of ProteinMPNN,^22^ with the addition of Gaussian noise (std-dev 0.14 Å) on the atomic coordinates of the proteins in the training set.

Effie was applied to the positive and negative states and the resulting decomposable score function was simplified based on the symmetrical properties of each state. The approximate Pareto front was produced for each design region, as described in Section. This produced 450 (resp. 778) solutions for the small (resp. large) region (see Figure 6). Following sequence clustering, the sequence minimizing the absolute value of the difference in Effie-score between the two negative states, 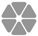 and 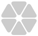, was selected in each cluster, producing 210 and 204 sequences for evaluation. To favor optimization of the positive state, only sequences for which |*λ*^−^| *< λ*^+^ were kept resulting in a final count of 189 (resp. 188) sequences.

**Figure 6:**
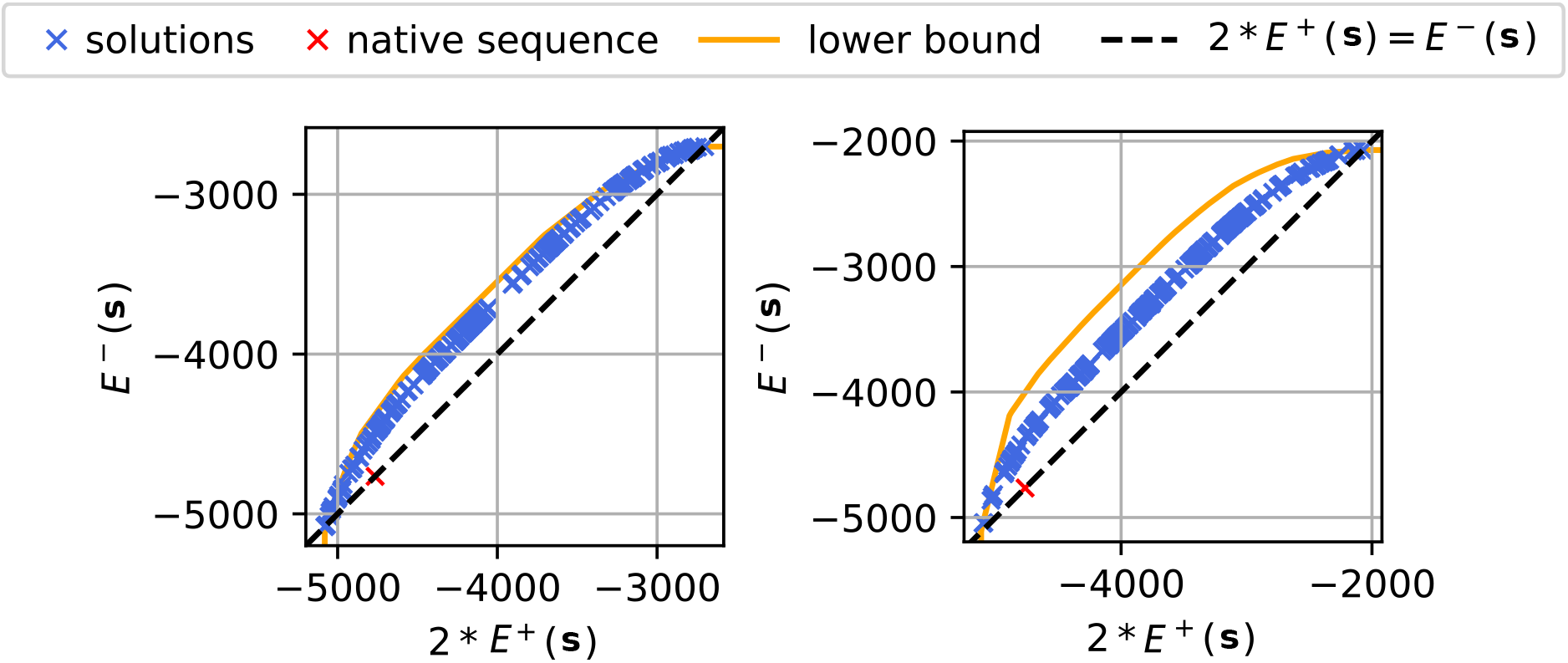
Approximate Pareto fronts proceed by Effie^*D*^ for the small (left) and large (right) regions.

#### ProteinMPNN-based design

ProteinMPNN was used with its default parameters: model `v 48 020` (trained with 0.20 Å Gaussian noise added on protein structures backbone), and a sampling temperature *T* = 0.1.

The positive state 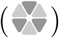 was assigned a weight *λ*^+^ = 1. As in the previous case, to give more importance to the positive state, weights *λ*^−^ ranging from −0.9 to −0.1 by increments of 0.1 were tested for the negative states (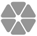and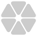), using the same weight for both states. Symmetrical chains in all states were tied using the `homooligomer` option of the `make pos neg tied positions dict.py` script of ProteinMPNN.

Combinations with a negative weight between −0.45 and −0.25 were determined to be the most suitable for our design objective. If *λ*^−^ *<* −0.45, the formation of hexamer 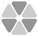 becomes energetically unfavorable. On the other hand, if *λ*^−^ is larger than −0.25, hexamers 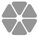 and 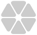 become favorable. This range of weights was explored with a finer resolution of 0.01 and for each weight combination, 10 designs were generated, for a total of 280 pairs of sequences for each designable region. Duplicate sequences were removed, leading to a total of 236 (resp. 259) designs.

#### Construction of mutant conformations

The designed sequences were mapped on the WT RMM symmetric hexamer conformation using our in-house Custozyme+ tool with a side chain packing of all residues using Rosetta `beta nov16` scoring function^63^ to create mutant conformations of the three different states considered in our multistate designs (one heterohexamer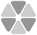 and two homohexamers 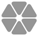 and 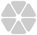). Mutant conformations were then minimized using the Rosetta minimize executable with the multi-step L-BFGS algorithm (`lbfgs armijo nonmonotone`), a minimization tolerance of 0.001 and `beta nov16` scoring function. No symmetry constraints were applied, allowing for large conformational change in the hexameric conformations. The resulting conformations are referred to as post-minimization conformations. Conformations pre- and post-minimization were scored using both Effie^*S*^ and ProteinMPNN^*S*^.

### Production and characterization of designed proteins

#### Cloning

Full-length DNA fragments coding for either Duo-GFP10 or Duo-GFP11 were synthesized by Twist Bioscience (sequences provided in List S2). Each pair of fragments was then Gibson-assembled within a NdeI/SalI-treated pET26b-based vector-1 (Kan^R^) that also coded for an His_6_-tagged GFP1-9 under an independent T7-transcription control. Gibson assembly reactions were carried out with the NEBuilder Hifi DNA Assembly Master Mix (E2621, NEB) following supplier instructions. Reactions were transformed in TOP10 cells and plasmids recovered by standard purification procedures.^60^ Final vectors were verified by restriction and sequencing (Eurofins).

DNA constructs for co-expression studies of Duo pairs in fusion to Flag peptide (Asp-Tyr-Lys-Asp-Asp-Asp-Asp-Lys) or His_6_ tags, which sequences are given in List S3, were directly cloned by Twist Bioscience between BglII and XhoI sites of their pET24 (+) (Kan^R^) expression vector.

#### Tripartite GFP assay: monitoring cellular growth and fluorescence

Protein expression studies were performed after the transformation of chemically-competent BL21(DE3) *Escherichia coli* cells with corresponding plasmids, which was carried out following standard protocols. A pre-culture of each strain in Luria-Bertani broth (LB, 100 µL) media supplemented with kanamycin (40 µg/mL final conc.: LBK media) was grown for 6 to 8 h at 37 °C and 200 rpm orbital-shaking. Then, 2 µL were seeded into LBK (200 µL) supplemented with IPTG (10 µM), previously dispensed in a 96-well black plate with glass flat bottom (Greiner, reference 655892). Cultures were continued in a CLARIOstar Plus (BMG Labtech), at 37 °C with continuous shaking. Optical density (at 600 nm) and GFP fluorescence (excitation wavelength at 470 *±* 15 nm, emission at 515 *±* 20 nm) were recorded approximately every 10 min, for 15 h. Data were fitted to the following sigmoidal function using PRISM 6 (GraphPad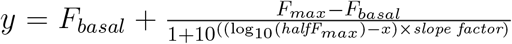 where *halfF*_*max*_ is the OD for which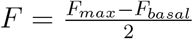.

For cases with too weak fluorescence readings, *F*_*max*_ values were extracted manually (typically the value read after 12 to 14 h of culture). Plotted averages and standard deviations derived from at least two independent experiments, each one including several replicates per case (typically 3 clones). Presented data were normalized with regard to the RMM reference value, since absolute fluorescence signal values varied substantially among independent experiments.

#### Protein expression, solubility and purification

The expression of proteins configured for the tripartite GFP assay (monomer A-GFP10 + monomer B-GFP11 + GFP1-9-His_6_), or consisting instead of A-Flag and B-His_6_ components (see above) was induced using same protocol. Namely, after growing 1 mL of pre-cultures overnight at 37 °C, 120 µL was inoculated in 12 mL of LBK. When the cultures reached mid-log phase (OD_600nm_ = 0.6 − 0.8), expression was induced with IPTG (200 µM final conc.). Incubation was continued for 4 hours before cell harvesting at 4,000 g, 4 °C. Supernatants were fully discarded and the pellets were gently resuspended in 0.8 mL of BugBuster (Novagen, amine free), supplemented with Benzonase (Proteogenix, 27 U/mL final) and lysozyme (20 µg/mL). The protease inhibitor phenylmethylsulfonyl fluoride was added immediately afterwards (PMSF, 1 mM final). After incubation at room temperature (RT) for 5 min, samples were transferred back to 4 °C before adding sodium phosphate (NaPi, 20 mM final, pH 8.1) and imidazole (10 mM). Total cellular content (TCC) fractions were then prepared from mixtures of 15 µL of these lysed samples and 45 µL of SDS loading dye (1.34X) and were denatured at 95 °C for 8 min. Insoluble debris and aggregates were removed by a 21,000 g centrifugation (10 min, 4 °C). Aliquots from resulting soluble fractions were denatured for sodium dodecylsulfate polyacrylamide gel electrophoresis (SDS-PAGE) analysis, as detailed for the TCC fraction. The remaining supernatant was loaded onto two Vivapure 8-96 well cobalt-chelate micro-columns (VivaScience) that had been preconditioned with Sol A (20 mM NaPi, 300 mM NaCl, 10 mM imidazole, pH 8.0). After 4 *×* 500 µL washing steps with Sol A, purified proteins were eluted with 300 µL of Sol B (Sol A plus 300 mM imidazole). EDTA was added (5 mM final conc.) and 60 µL of such purified fractions were denatured at 95 °C after mixing with 15 µL of SDS loading dye (5X).

Analysis by SDS–polyacrylamide gel electrophoresis was typically performed on 17 % gels. Loaded volumes of TCC (typically 4 µL), soluble (4 µL), and purified (6 µL) fractions were identical for all compared samples. Protein bands were visualized with Coomassie Brilliant Blue R250 (BioRad).

#### Western blot analysis

After SDS-PAGE, gel contents were electro-transferred to a PVDF membrane (Immobilion-P, Milipore). Membranes were blocked at RT for 1 h in 5 % non-fat dry milk resuspended in TBS containing 0.05 % Tween 20. Subsequent treatments and washing steps followed standard protocols. The primary antibody immunolabelling was carried out with a mouse monoclonal anti-Flag antibody (MA1-91878, ThermoFisher), diluted at 1:2000 in the blocking solution. After incubation with a similarly diluted alkaline phosphatase-conjugated goat anti-mouse IgG (H+L) secondary antibody (31346, Fisher Scientific), and washing steps, blots were developed by reaction with Sigmafast BCIP/NBT substrate (typically for 3 to 5 min).

#### Size-exclusion chromatography

Protein oligomeric state was estimated by SEC using a Beckman Ultraspherogel SEC2000 column (7.5 *×* 300 mm) operated by a Waters 2690 HPLC separation module. After equilibration of the column in 25 mM NaPi, 150 mM NaCl, pH 7.5, each protein sample (30 µL) was injected and run at 1 mL/min flow rate. Elution was monitored at 280 nm with a Waters 996 Photodiode Array Detector (absorption). Elution volumes permitted to estimate the species molecular weight by comparison to calibration standards run under identical con-ditions: dextran blue (2 MDa), ferritin (440 kDa), aldolase (158 kDa), conalbumin (75 kDa), ovalbumin (43 kDa), carbonic anhydrase (35 kDa) and ribonuclease A (13.7 kDa).

Separation of Duo4 oligomeric species was carried out in a Superdex 200 Increase 10/300 GL column, operated by an Akta Pure Protein Purification System. About 300 µL of the corresponding cobalt-chelate-purified sample was injected and run in the column preconditioned with 10 mM Tris HCl, 200 mM NaCl, 0.5 mM EDTA, at pH 7.8. Protein-containing peaks were collected manually (280 nm detection). Fractions corresponding to the same peak were pooled together and concentrated 20 to 40-fold using Vivaspin 500 concentrator devices (10 kDa MWCO, PES). Resulting samples were reanalyzed by SEC-HPLC as indicated above.

## Acknowledgement

This work was supported by the national research agency ANR (ANR-19-CE09-0032-02). The authors thank the regional computing meso center CALMIP for granting access to High Performance Computing resources.

## Supporting Information Available

Detailed methods of bi-objective optimization for negative design. Representation of the small and large designable regions on one interface between two WT RMM subunits. Comparisons of Effie^*S*^ and ProteinMPNN^*S*^ scores for pairs of sequences predicted on the large designable region. RMSD of the hexamer backbone as a function of their score computed either with Effie^*S*^ or ProteinMPNN^*S*^ for sequence pairs predicted on the small designable region. Comparison of Effie^*S*^ inter-chain scores for mutant sequences predicted either by Effie^*D*^ or ProteinMPNN^*D*^, on the small or large designable region. Experimental screening of the Duo designs using the tripartite GFP technology and the co-purification of partners with His-Tag GFP by SDS-PAGE. Characterization of the oligomeric state of Duo4 by HPLC and FPLC experiments (PDF). Sequences of 24 experimentally tested designs. Link to the Git repository for the toulbar2 bi-objective extension.

## TOC Graphic

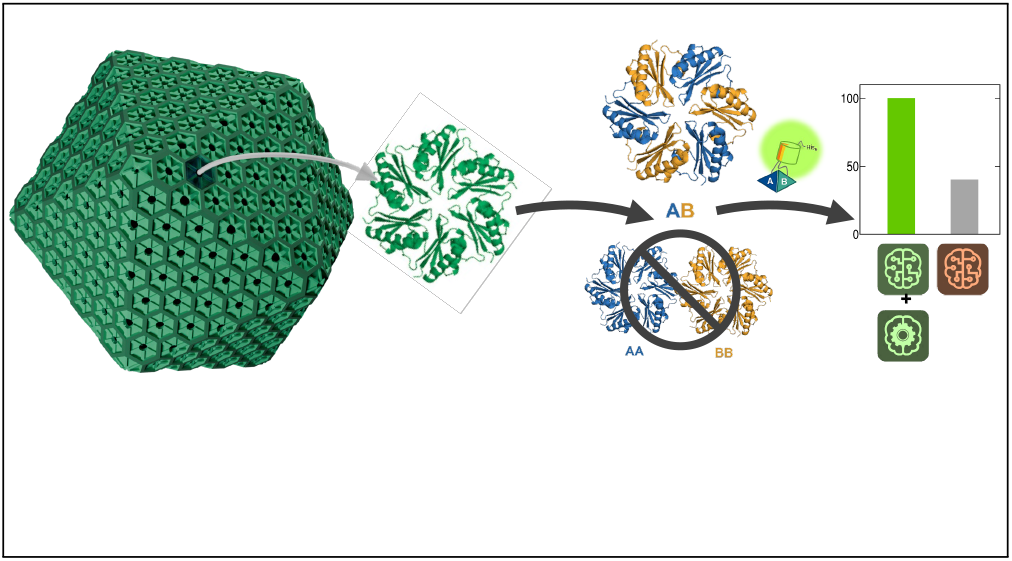

