## Supplemental Material for "Designing symmetrical multi-component proteins using a hybrid generative AI approach"

#### Bi-objective optimization for negative design

##### Definitions and properties

In this part, we recall several definitions and properties of bi-objective optimization problems. The sets $\mathbf{B}^+$ and $\mathbf{B}^-$ of positive and negative states define two criteria $E^+(\mathbf{s}) = \sum_{B_j \in \mathbf{B}^+} E(\mathbf{s}|B_j)$ and $E^-(\mathbf{s}) = \sum_{B_j \in \mathbf{B}^-} E(\mathbf{s}|B_j)$ which are the scores of the sequence $\mathbf{s}$ on these two sets of states.

Our multi-state computational protein design problem, based on the pairwise decomposable score function predicted by Effie, is formulated as a bi-objective combinatorial optimization problem:

$$\begin{aligned} &\underset{\mathbf{s}}{min} \quad E^{+}(\mathbf{s}) \\ &\underset{\mathbf{s}}{max} \quad E^{-}(\mathbf{s}) \end{aligned} \tag{1}$$

The score-pairs $(E^{+}(\boldsymbol{s}), E^{-}(\boldsymbol{s}))$ of various sequences can be compared through the notion of dominance, of efficient solutions and Pareto front, as defined below:

**Definition 1** (Dominance)**.** *A given solution (sequence)* $\boldsymbol{s}$ *is said to dominate another solution* ***s'*** *if and only if either* $E^{+}(\mathbf{s}) < E^{+}(\mathbf{s}')$ *and* $E^{-}(\mathbf{s}) \geq E^{-}(\mathbf{s}')$*, or* $E^{+}(\mathbf{s}) \leq E^{+}(\mathbf{s}')$ *and* $E^{-}(\mathbf{s}) > E^{-}(\mathbf{s}')$

**Definition 2** (Efficient solutions and Pareto front)**.** *A solution (sequence)* $\mathbf{s}$ *is called efficient for problem (1) if there exists no solution* $\mathbf{s}'$ *such that* $\mathbf{s}$ *dominates* $\mathbf{s}'$*. The Pareto front of the problem (1) is the set of score-pairs* $(E^{+}(\mathbf{s}), E^{-}(\mathbf{s}))$ *for all efficient sequences* $\mathbf{s}$*.*

Fig. s1* shows an example of a Pareto front (score-pairs of efficient solutions) for our *min* and *max* negative design problem. This example highlights the conflict between the two objectives: improving the score of one objective leads to the degradation of the other. In this figure, supported solutions are efficient solutions located on the convex hull of the Pareto front.

The linear scalarization of the problem merges the two objectives into one:

$$\underset{\mathbf{s}}{min} \quad \lambda^{+}E^{+}(\mathbf{s}) + \lambda^{-}E^{-}(\mathbf{s}) \tag{2}$$

With $\lambda^{+} \geq 0$ and $\lambda^{-} \leq 0$ since the first objective is to be minimized and the second objective is to be maximized. The following property shows that any solution that is optimal for the scalarization (2) is an efficient solution of the bi-objective problem 1.

**Theorem 1.** *Let* $\mathbf{s}$ *be an optimal solution of problem (2) for some* $\lambda^{+} \geq 0, \lambda^{-} \leq 0$*. Then* $\mathbf{s}$ *is an efficient (non-dominated) solution of problem (1).*

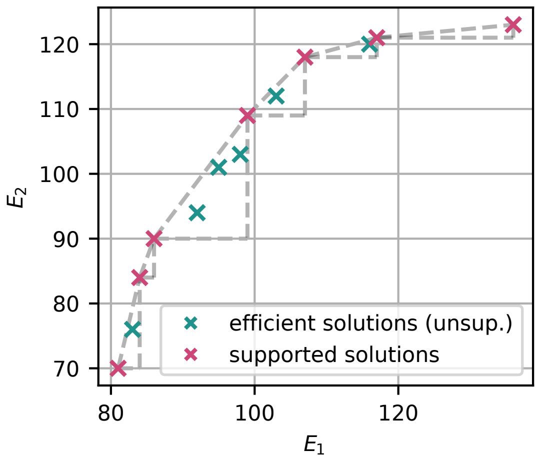


**Figure s1*:** Example of a Pareto front for our (min, max) negative design problem. Supported (resp. unsupported) efficient solutions are in red (resp. green).

*Proof.* Let $\mathbf{s}$ be an optimal solution of problem (2) and let us assume that there exists a solution $\mathbf{s}'$ which dominates $\mathbf{s}$ in problem (1). W.l.o.g. let us assume that $E^+(\mathbf{s}') < E^+(\mathbf{s})$ and $E^-(\mathbf{s}') = E^-(\mathbf{s})$. This implies $\lambda^+ E^+(\mathbf{s}') + \lambda^- E^-(\mathbf{s}') = \lambda^+ E^+(\mathbf{s}') + \lambda^- E^-(\mathbf{s}) < \lambda^+ E^+(\mathbf{s}) + \lambda^- E^-(\mathbf{s})$ This would imply that $\mathbf{s}$ is not optimal for objective (2), which is a contradiction. □

Additionally, solutions that are optimal for the scalarization problem are also called supported solutions of the bi-objective problem in the sense that their score-pairs are located on the convex hull of the Pareto front (see Fig. s1*).

#### Approximation Dichotomy Algorithm

Our approach to solve problem (1) follows a two-phase approximation method where the goal of the second phase is to produce a dense set of solutions (without Pareto optimality guarantee):

1 $Dichotomy(N, t, m, HBFS)$ produces an initial set of solutions as well as a dual bound of the Pareto front ; it returns a list of solutions and a list of pairs of weights associated with each dual bound $(L)$ ;

2 $Dichotomy(N, t, m, PerturbedVNS(nbrepeat, \epsilon))$ produces a set of varied solutions using a metaheuristic, Variable Neighborhood Search (VNS).[S1] To find a dense set of solutions, VNS is restarted *nbrepeat* times with a perturbed objective function (with the same scalarization weights) obtained by adding a small random uniform noise (in $[0, \epsilon]$) on the scalarized objective. Only a primal bound of the Pareto front is returned ($U$).

We used $nbrepeat = 5$, $\epsilon = 0.005$.

Our approximation dichotomy search algorithm (items 1 and 2 above) is further described in algorithm 1. It relies on the call to the function $Solve(\lambda^+, \lambda^-, ub, t)$ which solves a linear scalarization of problem 1. This function takes as input a pair of weights $(\lambda^+, \lambda^-)$ for the scalarization, an upper bound $ub$ (sequences with score greater than $ub$ are not sought), and a time limit $t$. It returns a tuple $(S, l)$ where $l$ is a lower bound on the optimum of the scalarization and $S$ is a set of solutions found during search (multiple runs with perturbations can be performed, returning several solutions). The approximation dichotomy algorithm has been implemented with the Cost Function Network solver toulbar2,[S2] (https://github.com/toulbar2/toulbar2) relying on two algorithms within it: *Hybrid Best First Search (HBFS)*[S3] was used for exact search, Variable Neighborhood Search (VNS)[S1] was used as the metaheuristic. The source code of this bi-objective extension is available at https://forgemia.inra.fr/samuel.buchet/boond.

In the main loop of the algorithm, the construction of a new pair of weights is modified from the original Dichotomy method[S4] through the use of the function *GetCandidate* as illustrated in Fig. s2*. This function allows for the control of the progression of the Pareto front approximation as well as the number of iterations of the algorithm when the output solutions of *Solve* are not proven to be optimal.

**Function** Dichotomy($N, t, m, Solve$)

$L := \varnothing$; $U := \varnothing$; $Q := \varnothing$; $W := \varnothing$;

/* Solve $1. * E^{+}(\boldsymbol{s}) + 0. * E^{-}(\boldsymbol{s})$ within time $t$, store upper & lower bound */

$S_1, l_1 :=$ Solve$(1., 0., \infty, t)$ ; $\boldsymbol{x_1^*} :=$ pop-first$(S_1)$ ;

$L$.push$(\langle 1., 0., l_1 \rangle)$;

$U$.merge$(S_1)$;

/* Solve $0. * E^{+}(\boldsymbol{s}) - 1. * E^{-}(\boldsymbol{s})$ within time $t$, store upper & lower bound */

$S_2, l_2 :=$ Solve$(0., -1., \infty, t)$ ; $\boldsymbol{x_2^*} :=$ pop-first$(S_2)$;

$L$.push$(\langle 0., -1., l_2 \rangle)$;

$U$.merge$(S_2)$;

/* Proceed if $F_1$ and $F_2$ have been solved to optimality */

**if** $\boldsymbol{x_1^*} \neq \varnothing \wedge \boldsymbol{x_2^*} \neq \varnothing$ **then**

  $Q := \{(\boldsymbol{x_1^*}, \boldsymbol{x_2^*})\}$

**while** $Q \neq \varnothing$ **and** $|L_1| \leq m$ **do**

  /* Bisect by $(\lambda_1, \lambda_2)$-scalarization and store bounds */

  $(\boldsymbol{x_1}, \boldsymbol{x_2}) :=$ pop-first$(Q)$;

  $\lambda^{+} := E^{-}(\boldsymbol{x_1}) - E^{-}(\boldsymbol{x_2})$; $\lambda^{-} := E^{+}(\boldsymbol{x_2}) - E^{+}(\boldsymbol{x_1})$;

  $S, l :=$ Solve$(\lambda^{+}, \lambda^{-}, \lambda^{+}E^{+}(\boldsymbol{x_1}) + \lambda^{-}E^{-}(\boldsymbol{x_2}), t)$;

  $\boldsymbol{x} :=$GetCandidate$(S, \boldsymbol{x_1}, \boldsymbol{x_2}, \lambda^{+}, \lambda^{-})$ ;

  $L$.push$(\langle \lambda^{+}, \lambda^{-}, l \rangle)$ ;

  $U$.merge$(S)$ ;

  /* Push $\boldsymbol{x}$ in $Q$ only if it improves the Pareto frontier approximation */

  **if** $\boldsymbol{x} \neq \varnothing$ **then**

    $Q$.push$((\boldsymbol{x_1}, \boldsymbol{x}))$ ; $Q$.push$((\boldsymbol{x}, \boldsymbol{x_2}))$ ;

  **else**

    $W$.push$((\boldsymbol{x_1}, \boldsymbol{x_2}))$

**return** $L, U, W$

**Function** GetCandidate($S, \boldsymbol{x_1}, \boldsymbol{x_2}, \lambda^{+}, \lambda^{-}$)

$\boldsymbol{x_{new}} := \emptyset$;

**for** $\boldsymbol{x} \in S$ **do**

  $select := True$

  **if** $\boldsymbol{x_{new}} \neq \emptyset \wedge \lambda^{+}E^{+}(\boldsymbol{x}) + \lambda^{-}E^{-}(\boldsymbol{x}) \geq \lambda^{+}E^{+}(\boldsymbol{x_{new}}) + \lambda^{-}E^{-}(\boldsymbol{x_{new}})$ **then**

    $select := False$

  **if** $\vec{v_1} \cdot \vec{v_2} < 0 \vee \vec{v_1} \cdot \vec{v_3} < 0$ **then**

    $select := False$

  $\vec{v_1} := \begin{pmatrix} -\lambda^{+} \\ -\lambda^{-} \end{pmatrix}$ ; $\vec{v_2} := \begin{pmatrix} E^{+}(\boldsymbol{x}) - E^{+}(\boldsymbol{x_1}) \\ E^{-}(\boldsymbol{x}) - E^{-}(\boldsymbol{x_1}) \end{pmatrix}$ ; $\vec{v_3} := \begin{pmatrix} E^{+}(\boldsymbol{x}) - E^{+}(\boldsymbol{x_2}) \\ E^{-}(\boldsymbol{x}) - E^{-}(\boldsymbol{x_2}) \end{pmatrix}$

  **if** $\vec{v_1} \cdot \vec{v_2} < 0 \vee \vec{v_1} \cdot \vec{v_3} < 0$ **then**

    $select := False$

  **if** $select$ **then**

    $\boldsymbol{x_{new}} := \boldsymbol{x}$

**return** $\boldsymbol{x_{new}}$

**Algorithm 1:** Approximation dichotomy algorithm to approximate a Pareto front.

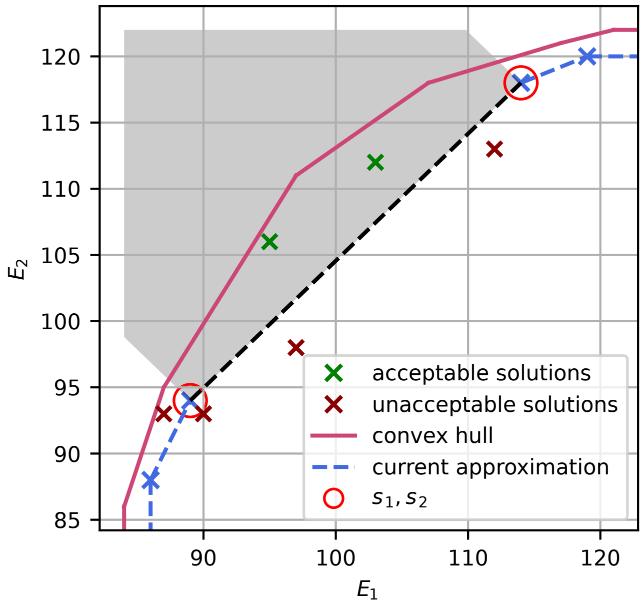


**Figure s2*:** Representation of the selection of a candidate solution (function *GetCandidate*) to form new pairs of weights in our modified Dichotomy algorithm. The green and red crosses represent solutions returned by a call to function *Solve*. The grey area represents the area where solutions can be considered to form new pairs of weights. In this area, the farthest solution from the black line (i.e. with the best scalarized score) is to be selected to produce two new pairs of weights by association with the input solutions $\boldsymbol{s_1}$ and $\boldsymbol{s_2}$.

#### Comparison of the two phases

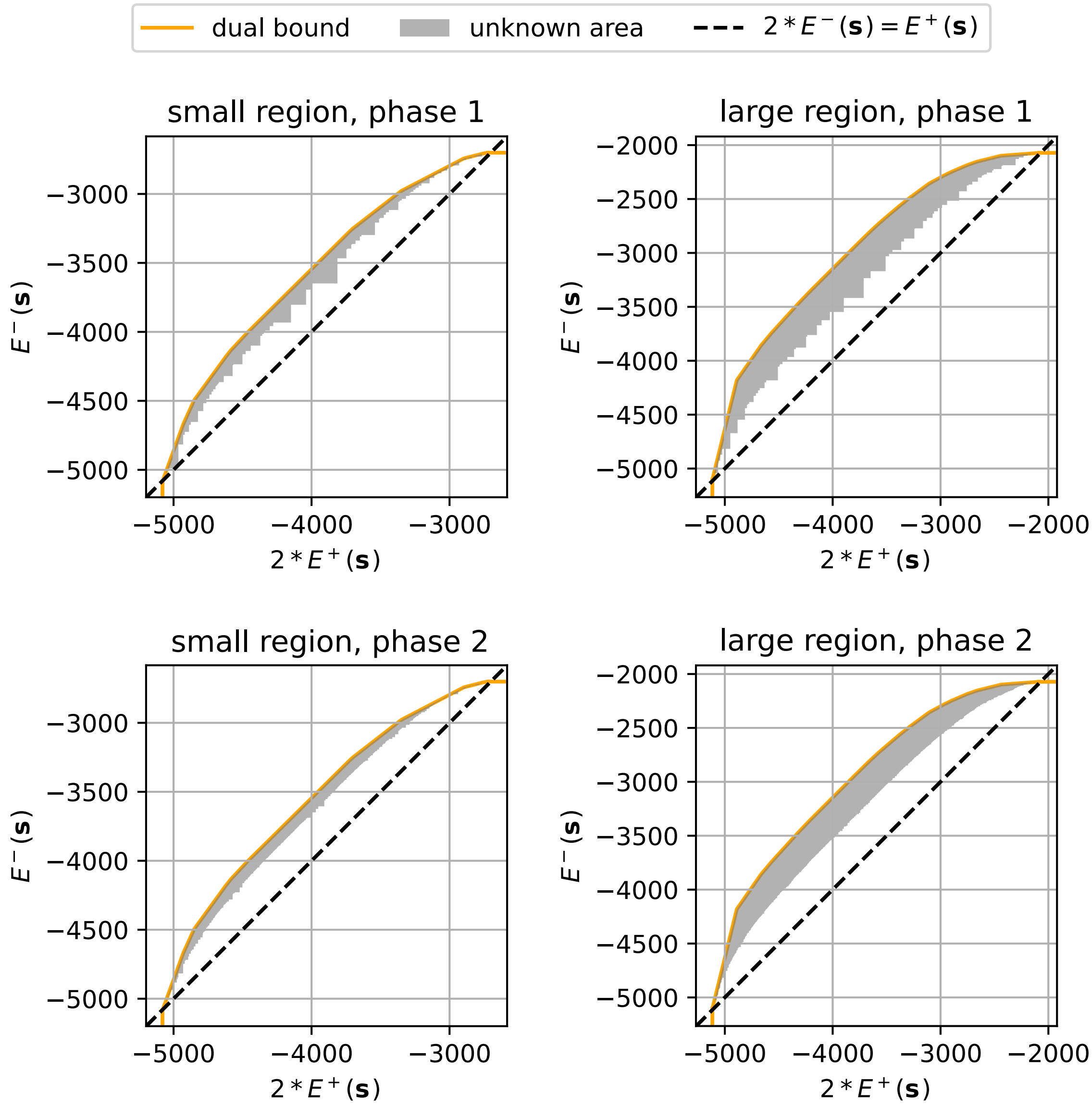


**Figure s3*:** Improvement of the approximation of the Pareto front with the second phase of the method on each mutable region.

#### Comparison of the two designable regions

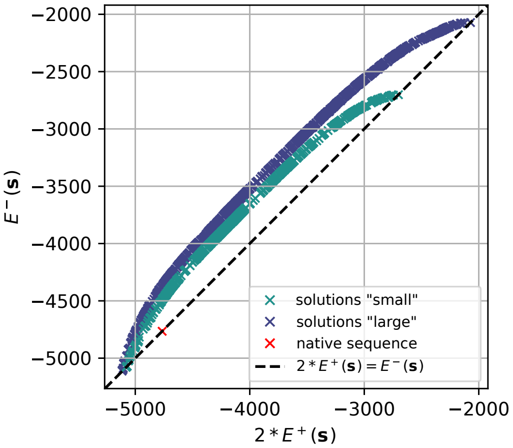


**Figure s4*:** Superposition of the score-pairs of the Pareto fronts approximations computed for the two designable regions. Since the larger region includes the residue positions of the smaller ones, the solutions for that region dominate the other ones, with a shift toward higher values of the Effie$^S$ score.

### Design states and designable region

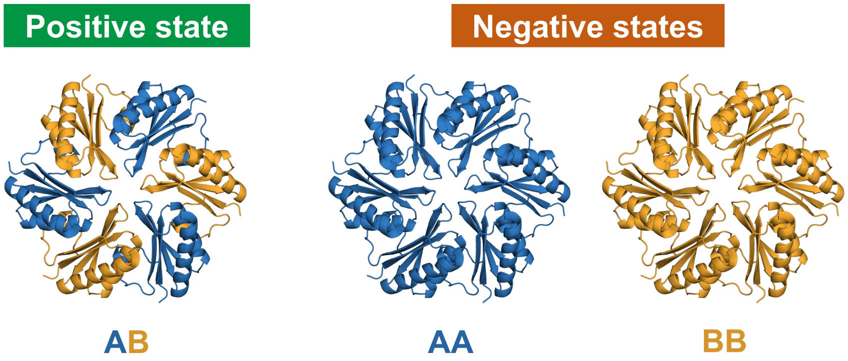


**Figure S1:** Representation of the positive state, heterohexamer AB, and both negative states, homohexamer AA and BB, used in our design approaches from the RMM WT symmetric template conformation reconstructed using Rosetta C6 symmetry with ref2015 score function.

.

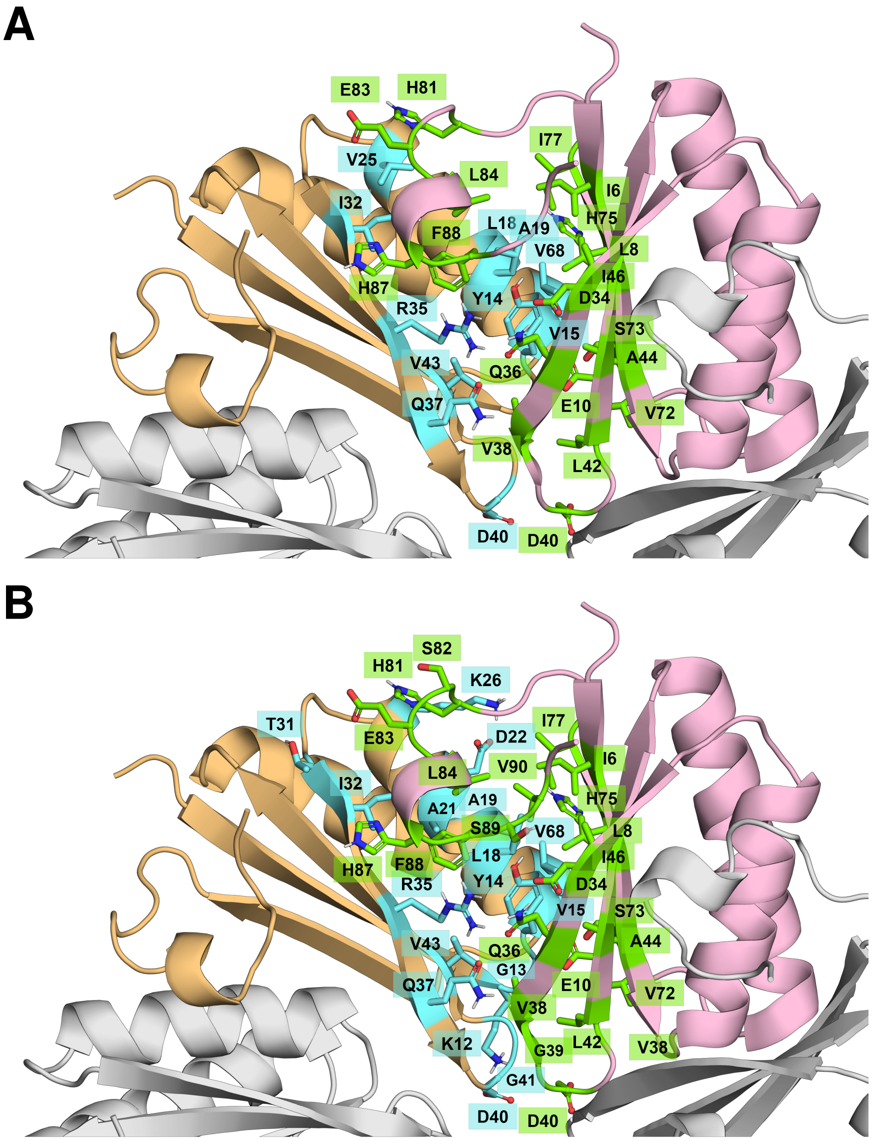


**Figure S2:** Interface between two monomers from WT RMM with the designable residues from the small (**A**) or the large region (**B**). The hexamer backbone is showed in cartoon representation with two subunits involved in one interface colored either in orange or in pink. Each side of the interface is colored either in cyan or in green with the side chains' atoms colored by type. For the small region (**A**), the first side of the interface is composed of **Y14, V15, L18, A19, V25, I32, R35, Q37, D40, V43 and V68**, and the second side of the interface of **I6, L8, E10, D34, Q36, V38, D40, L42, A44, I46, V72, S73, H75, I77, H81, E83, L84, H87 and F88**. For the large region (**B**), the first side of interface is composed of **K12, G13, Y14, V15, L18, A19, A21, D22, K26, T31, I32, R35, Q37, D40, G41, V43 and V68**, and the second side of the interface of **I6, L8, E10, D34, Q36, V38, G39, D40, L42, A44, I46, V72, S73, H75, I77, H81, S82, E83, L84, H87, F88, S89 and V90**.

### Comparison between Effie and ProteinMPNN solutions

#### Figures

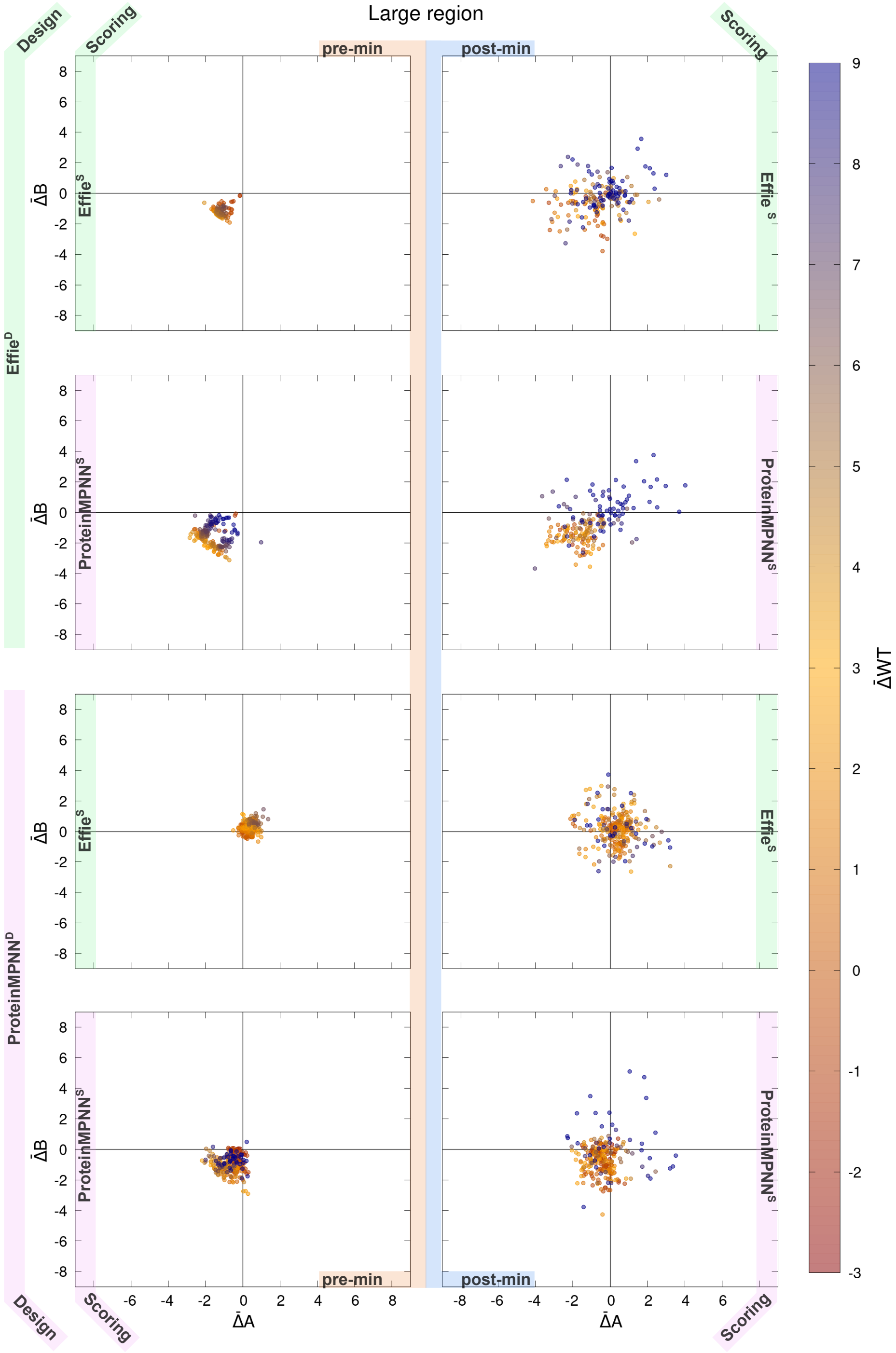


**Figure S3:** Comparison of scores for mutants generated by Effie$^D$ or ProteinMPNN$^D$ on the large designable region.

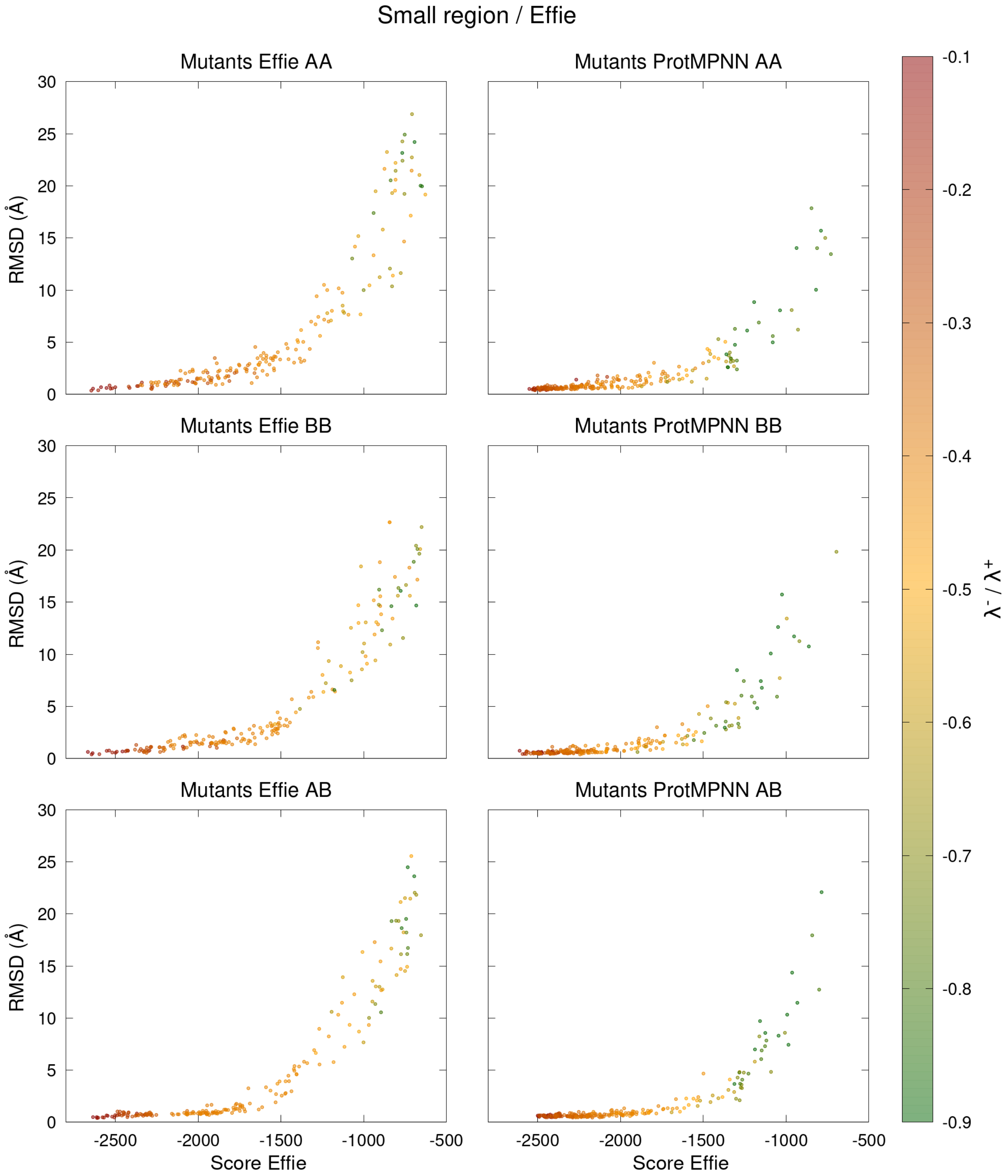


**Figure S4:** RMSD of the hexamer backbone of each state as a function of their Effie$^S$ score for mutant sequences predicted on the small region by either Effie$^D$ (left) or ProteinMPNN$^D$ (right).

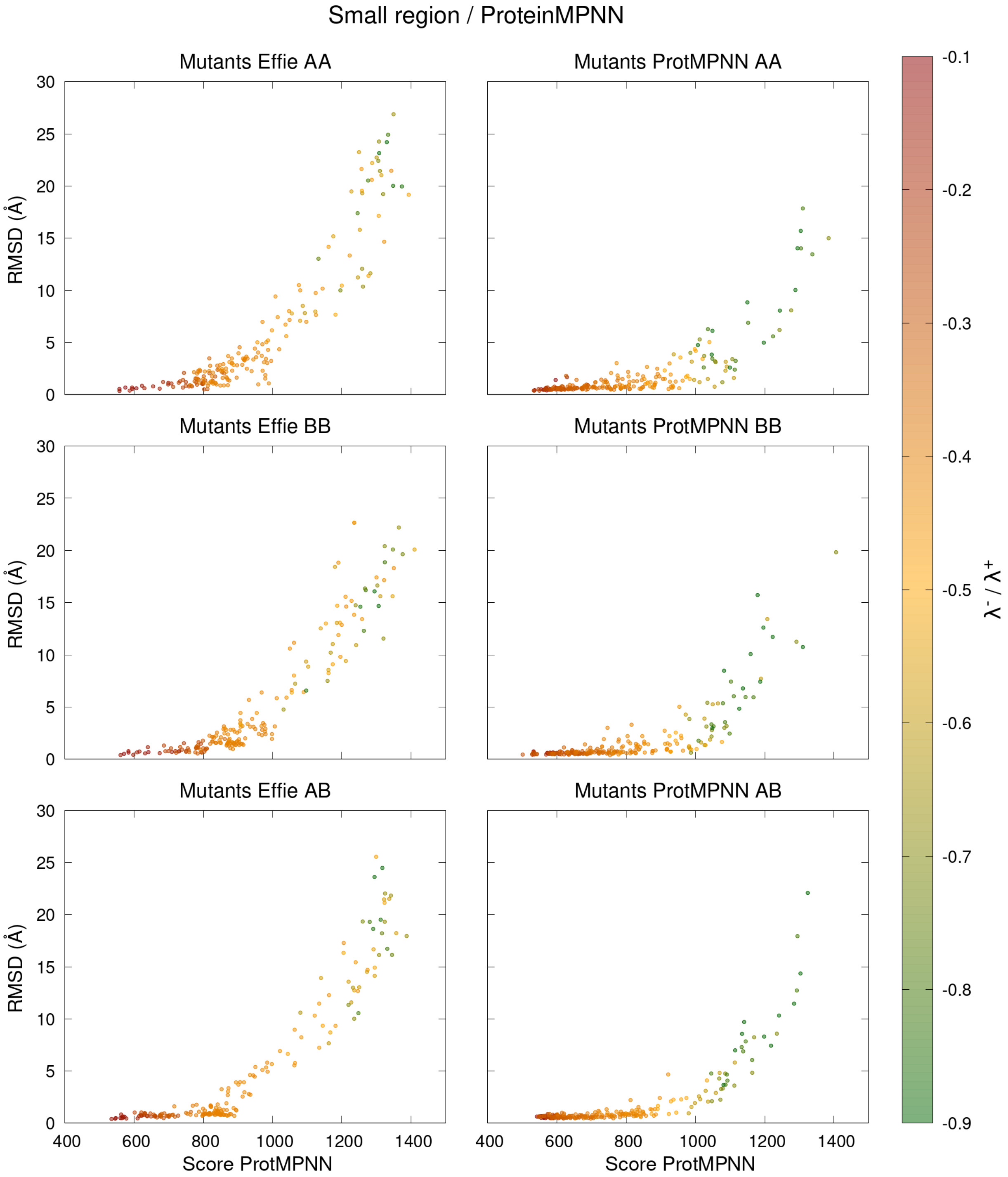


**Figure S5:** RMSD of the hexamer backbone of each state as a function of their ProteinMPNN$^S$ score for mutant sequences predicted on the small region by either Effie$^D$ (left) or ProteinMPNN$^D$ (right).

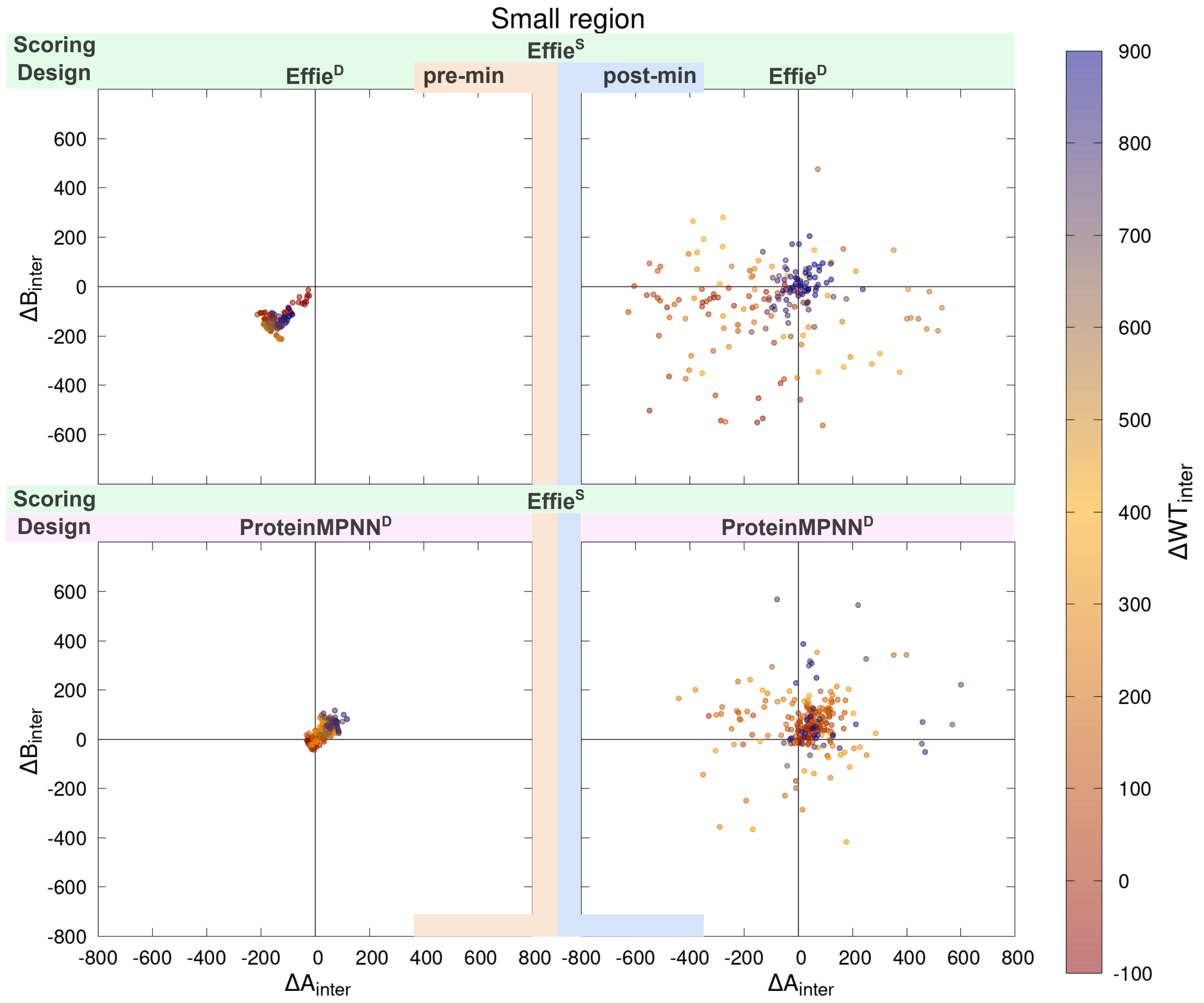


**Figure S6:** Comparison of Effie$^S$ inter-chain scores for mutant sequences predicted either by Effie$^D$ or ProteinMPNN$^D$ on the small designable region, pre- (left) or post-minimization (right).

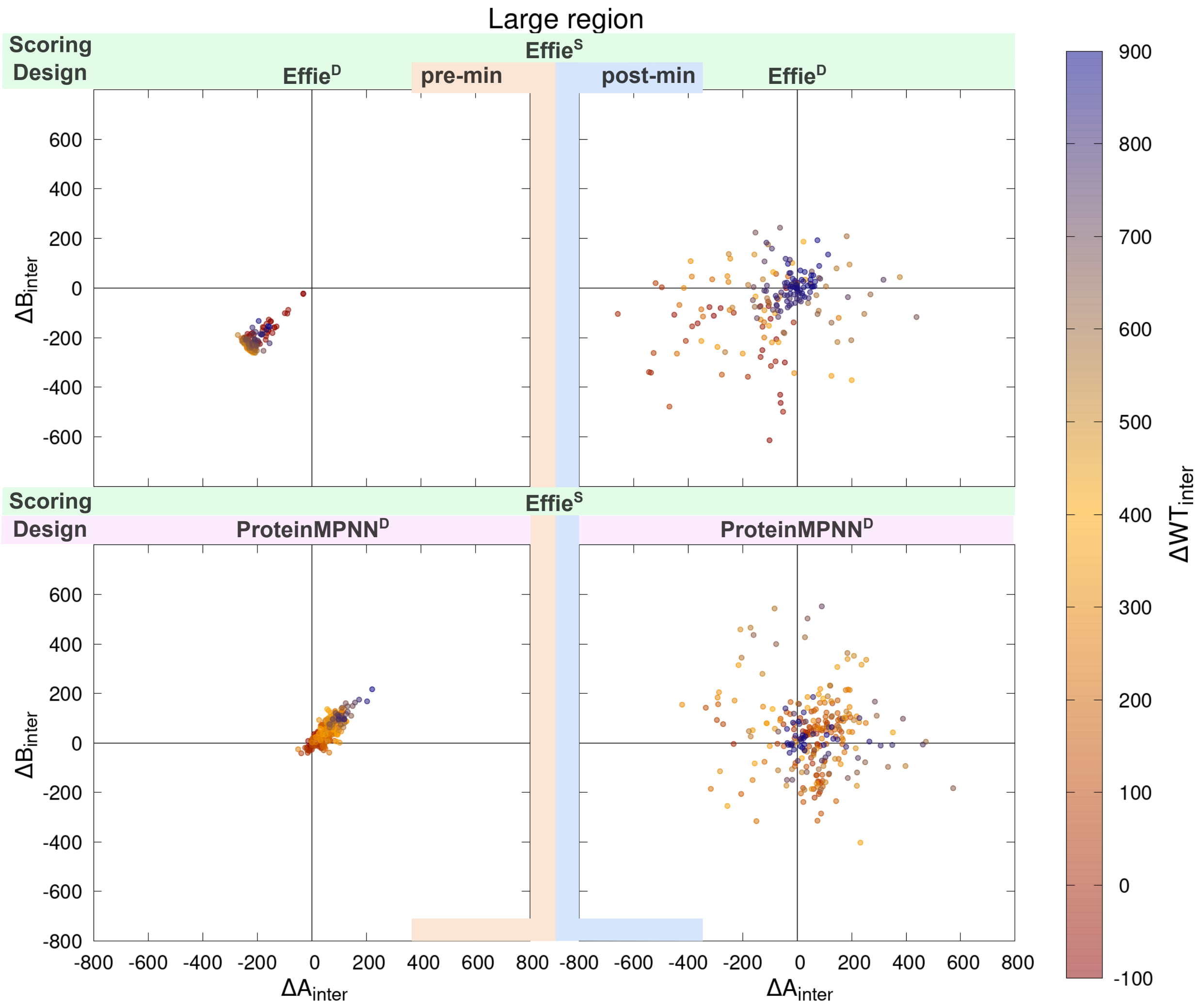


**Figure S7:** Comparison of Effie$^S$ inter-chain scores for mutant sequences predicted either by Effie$^D$ or ProteinMPNN$^D$ on the large designable region, pre- (left) or post-minimization (right).

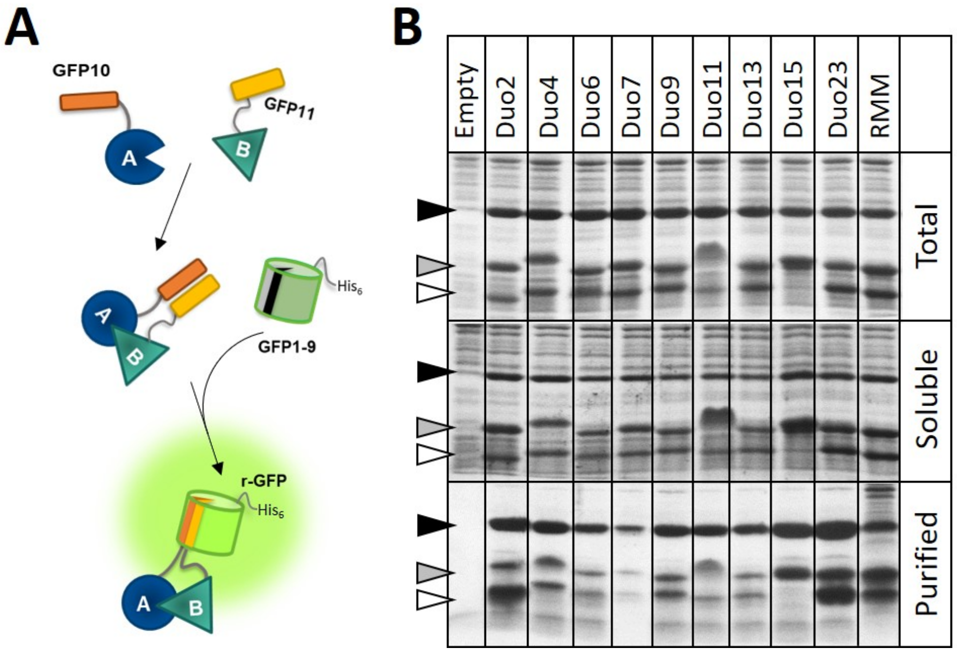


**Figure S8:** Experimental screening of the Duo designs using the tripartite GFP technology. (**A**) Schematic explanation of the tripartite GFP technology. An adapted GFP, composed of 11 $\beta$-strands, is split in 3 parts: the last two strands, GFP10 and GFP11, plus the remaining portion, GFP1-9. The small strands are connected to proteins A and B being screened. For an interacting A/B pair, GFP10 and GFP11 are brought in close proximity, thus facilitating the reconstitution of a full GFP (r-GFP). On the contrary, when proteins A and B are not good interactors, the GFP reconstitution (encounter of the three parts) is entropically hampered. (**B**) Purification of selected Duo designs. Cells overexpressing each designed pair of A-GFP10 and B-GFP11 monomers, plus the $His_6$-tagged GFP1-9 component, were collected at the end of the culture. Corresponding total cellular contents were analyzed in Coomassie blue-stained polyacrylamide gels (Top). Fractions remaining soluble after lysis and centrifugation (middle) were loaded on cobalt-based chelating resins, with the intention to retain $His_6$-tagged GFP1-9 and any bound partner. The resulting purified fractions are shown in the bottom of the panel. Further details are given in Figure 3 (see also Materials and Methods).

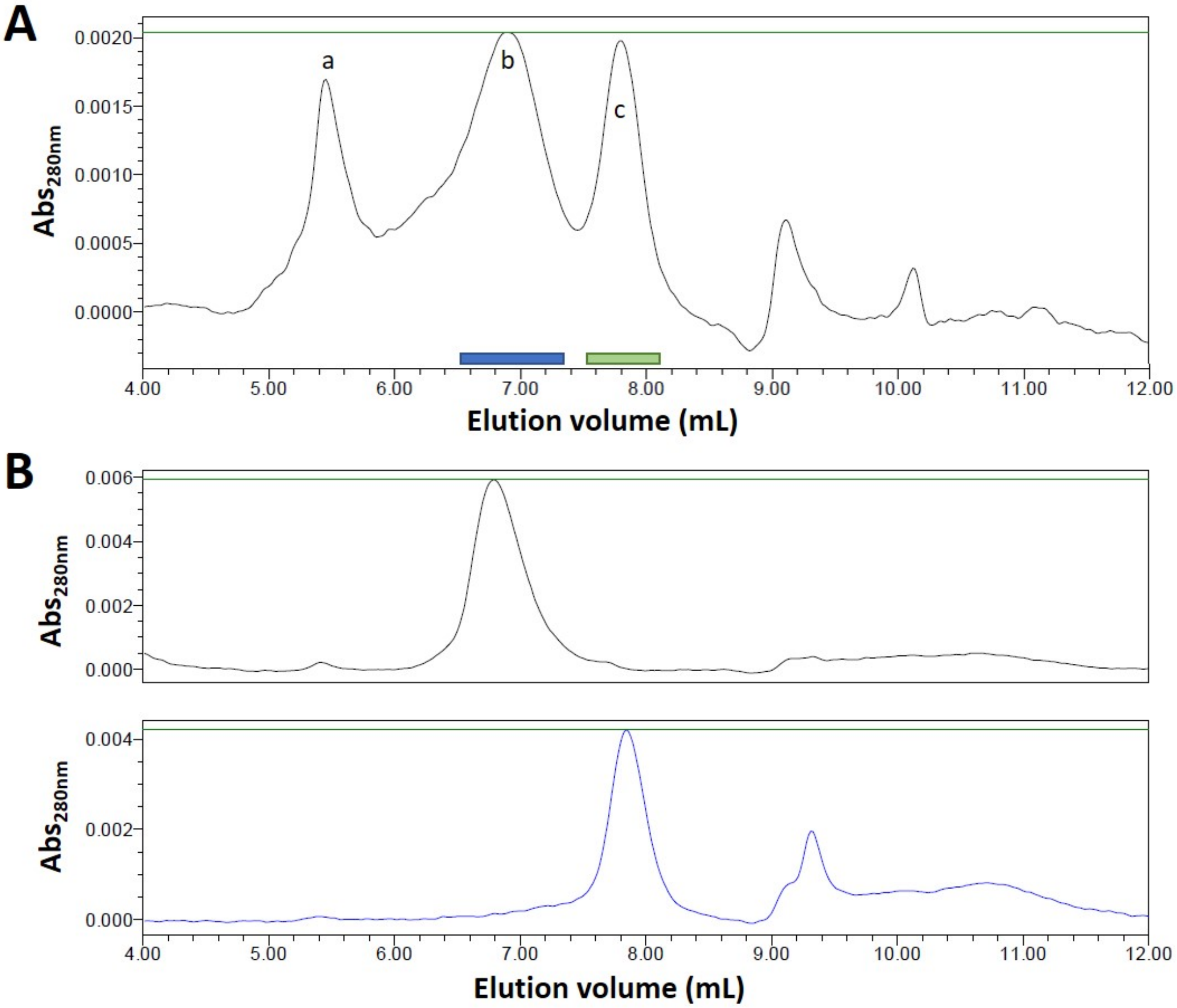


**Figure S9:** Characterization of the oligomeric state of Duo4. **A.** Size-exclusion chromatogram after injection of purified Duo4 on an analytical Beckman S2000 HPLC column. Shown is the profile of Duo4 absorption at 280 nm. Approximate molecular weights of 70 and 23 kDa were estimated for species eluting within peaks *b* and *c*, whereas the peak *a* likely consist of aggregated material (> 2 MDa). Elution at volumes higher than 9 mL would correspond to proteolysis (< 5 kDa) and/or small buffer components. Please refer to Materials and Methods for further details. **B.** Stability of isolated species displaying different oligomeric states. The purified Duo4 was injected in a Superdex 200 10/300 column (FPLC mode). This permitted to collect peaks *b* and *c* separately (the different fractions collected are indicated in panel A by blue and green bars). These fractions were concentrated and analyzed exactly as in panel A (on top for peak *a*, bottom for peak *b*).

#### Tables

**Table S1:** Evaluation of inter-chain Effie$^S$ scores for Effie$^D$ or ProteinMNN$^D$ designs for each designable region. ($n$) is the number of designs generated, ($\gamma_{\text{inter}}$) indicates how much the positive state is improved over the negative states when considering only Effie inter-chain scores, (yield) gives the percentage of designs with both $\bar{\Delta}A_{\text{inter}} < 0$ and $\bar{\Delta}B_{\text{inter}} < 0$.

| Scoring function | → | | Effie$^S$ | | | |
|---|---|---|---|---|---|---|
| | | | pre-min | | post-min | |
| Design function | Design space | $n$ | $\gamma_{\text{inter}}$ | yield | $\gamma_{\text{inter}}$ | yield |
| Effie$^D$ | small | 189 | 1.74 | 100 % | 0.87 | 43.4 % |
| | large | 188 | 2.52 | 100 % | 0.77 | 45.7 % |
| PMPNN$^D$ | small | 236 | -0.40 | 6.4 % | -0.59 | 8.1 % |
| | large | 259 | -0.68 | 4.6 % | -0.54 | 10.4 % |

**Table S2:** Evaluation by Effie$^S$ of the pairs of mutant sequences generated using either Effie$^D$ or ProteinMPNN$^D$ selected for experimental characterization.

| Scoring function → | Effie$^S$ | | | | | | |
|---|---|---|---|---|---|---|---|
| Design function | pre-min* | | | post-min | | | Mutant name |
| | $\Delta A$ | $\Delta B$ | $\Delta WT$ | $\Delta A$ | $\Delta B$ | $\Delta WT$ | |
| Effie$^D$ | -32.3 | -149.6 | 170.4 | -80.4 | -161.1 | 223.2 | Duo1 |
| | -60.4 | -171.1 | 70.6 | -223.4 | -336.1 | 179.4 | Duo2 |
| | -157.0 | -59.4 | 201.6 | -813.5 | -163.2 | 296.7 | Duo3 |
| | -73.4 | -86.6 | 139.6 | -107.6 | -592.3 | 223.2 | Duo4 |
| | -146.4 | -170.5 | 144.9 | -611.9 | -210.9 | 198.7 | Duo5 |
| | -59.1 | -60.6 | 112.1 | 77.9 | -300.3 | 130.4 | Duo6 |
| | -49.6 | -128.8 | 60.1 | -201.6 | -204.4 | 43.1 | Duo7 |
| | -103.2 | -147.8 | 35.2 | -226.5 | -327.4 | 97.9 | Duo8 |
| | 4.9 | -138.5 | 111.3 | -103.6 | -86.0 | 226.5 | Duo9 |
| | -30.3 | -133.7 | 158.2 | -138.7 | -24.2 | 303.2 | Duo10 |
| | -76.4 | -52.6 | 194.0 | -134.7 | -292.1 | 285.3 | Duo11 |
| | -25.7 | -163.1 | 75.4 | -129.6 | -169.7 | 94.4 | Duo12 |
| | -51.6 | -104.9 | 192.0 | -444.8 | -44.3 | 114.3 | Duo13 |
| | -60.1 | -151.9 | 147.6 | -486.8 | -169.5 | 108.6 | Duo14 |
| ProteinMPNN$^D$ | 3.1 | 15.4 | 249.1 | -45.5 | -39.3 | 313.2 | Duo15 |
| | -30.1 | 55.7 | 134.6 | 120.3 | 51.0 | 172.1 | Duo16 |
| | 14.2 | 52.9 | 304.5 | -322.0 | 93.5 | 361.7 | Duo17 |
| | 31.7 | -5.7 | 270.0 | 35.8 | -297.6 | 258.9 | Duo18 |
| | 106.2 | 58.5 | 427.0 | 99.1 | -266.7 | 465.3 | Duo19 |
| | 93.1 | 42.0 | 495.2 | 81.2 | 232.7 | 849.3 | Duo20 |
| | 24.3 | 73.1 | 400.2 | -117.8 | -8.5 | 611.4 | Duo21 |
| | -32.0 | 11.3 | 272.4 | 231.7 | 176.8 | 693.1 | Duo22 |
| | -4.5 | -7.5 | 136.0 | -332.4 | -355.8 | 179.3 | Duo23 |
| | 38.8 | 51.0 | 210.2 | -189.5 | 100.3 | 208.1 | Duo24 |

* Mutant sequence pairs were scored on the WT template conformation.

**Table S3:** Evaluation by ProteinMPNN$^S$ of the pairs of mutant sequences generated using either Effie$^D$ or ProteinMPNN$^D$ selected for experimental characterization.

| Scoring function → | ProteinMPNN$^S$ | | | | | | |
|---|---|---|---|---|---|---|---|
| Design function | pre-min* | | | post-min | | | Mutant name |
| | $\Delta A$ | $\Delta B$ | $\Delta WT$ | $\Delta A$ | $\Delta B$ | $\Delta WT$ | |
| Effie$^D$ | -45.1 | -77.6 | 140.2 | -82.0 | -85.2 | 83.4 | Duo1 |
| | -53.2 | -91.2 | 103.1 | -60.7 | -59.5 | 89.3 | Duo2 |
| | -141.5 | -42.3 | 192.0 | -239.5 | -21.1 | 163.2 | Duo3 |
| | -21.6 | -71.5 | 153.9 | -37.1 | -144.4 | 52.1 | Duo4 |
| | -92.0 | -97.6 | 143.4 | -142.7 | -57.8 | 124.4 | Duo5 |
| | -45.6 | -64.7 | 111.4 | -59.8 | -105.8 | 36.0 | Duo6 |
| | -67.5 | -75.4 | 80.3 | -64.6 | -117.6 | 10.5 | Duo7 |
| | -98.8 | -97.2 | 76.4 | -96.2 | -69.9 | 30.2 | Duo8 |
| | -10.9 | -96.7 | 107.9 | 11.2 | -74.4 | 32.7 | Duo9 |
| | -43.0 | -94.9 | 154.9 | -61.1 | -65.8 | 46.4 | Duo10 |
| | -62.9 | -56.2 | 156.3 | -22.8 | -98.8 | 116.9 | Duo11 |
| | -43.0 | -109.6 | 94.2 | -9.9 | -126.1 | 28.4 | Duo12 |
| | -36.3 | -59.3 | 172.2 | -35.3 | -92.2 | 71.0 | Duo13 |
| | -44.0 | -86.0 | 137.0 | -72.6 | -87.3 | 76.0 | Duo14 |
| ProteinMPNN$^D$ | -39.2 | -32.9 | 125.3 | -46.4 | -73.9 | 92.5 | Duo15 |
| | -75.2 | -29.5 | 47.5 | -16.4 | -29.9 | 37.4 | Duo16 |
| | -65.7 | -45.9 | 105.1 | -76.3 | -44.0 | 104.3 | Duo17 |
| | -84.4 | -76.2 | 152.6 | -62.1 | -82.1 | 107.2 | Duo18 |
| | -89.8 | -68.5 | 202.0 | -67.2 | -92.2 | 133.8 | Duo19 |
| | -83.3 | -100.8 | 254.0 | -119.3 | -51.3 | 214.8 | Duo20 |
| | -69.0 | -115.4 | 213.9 | -57.6 | -56.6 | 194.3 | Duo21 |
| | -104.7 | -40.0 | 118.9 | -75.7 | -68.4 | 93.3 | Duo22 |
| | -59.8 | -79.2 | 22.5 | -95.1 | -142.8 | -26.8 | Duo23 |
| | -42.1 | -77.8 | 89.5 | -64.5 | -100.4 | 4.8 | Duo24 |

* Mutant sequence pairs were scored on the WT template conformation.

**Table S4:** Evaluation of the Effie$^S$ inter-chain scores for mutant sequence pairs generated using either Effie$^D$ or ProteinMPNN$^D$ selected for experimental characterization.

| Scoring function → | Effie$^S$ | | | | | | |
|---|---|---|---|---|---|---|---|
| | pre-min* | | | post-min | | | |
| Design function | $\Delta A_{inter}$ | $\Delta B_{inter}$ | $\Delta WT_{inter}$ | $\Delta A_{inter}$ | $\Delta B_{inter}$ | $\Delta WT_{inter}$ | Mutant name |
| Effie$^D$ | -74.2 | -107.1 | 2.6 | -27.6 | -95.9 | 123.1 | Duo1 |
| | -105.2 | -127.4 | -19.8 | -208.4 | -299.8 | 120.5 | Duo2 |
| | -114.1 | -102.1 | 4.2 | -696.6 | -232.1 | 117.7 | Duo3 |
| | -87.1 | -72.9 | 9.5 | -116.2 | -584.9 | 147.3 | Duo4 |
| | -166.9 | -150.0 | -33.8 | -577.4 | -225.4 | 54.8 | Duo5 |
| | -71.8 | -47.2 | -3.1 | 62.2 | -242.4 | 97.3 | Duo6 |
| | -88.1 | -90.3 | -22.6 | -181.2 | -147.2 | 29.3 | Duo7 |
| | -130.7 | -120.4 | -50.5 | -203.8 | -314.5 | 67.7 | Duo8 |
| | -55.2 | -78.5 | 16.4 | -101.3 | -43.2 | 248.5 | Duo9 |
| | -64.6 | -99.5 | 8.0 | -144.3 | -88.5 | 239.7 | Duo10 |
| | -65.6 | -63.8 | 33.8 | -213.9 | -247.0 | 164.1 | Duo11 |
| | -78.4 | -110.1 | -2.9 | -123.1 | -83.5 | 96.6 | Duo12 |
| | -71.1 | -85.5 | 28.3 | -408.1 | -34.8 | 25.7 | Duo13 |
| | -96.1 | -116.0 | -0.6 | -485.6 | -105.7 | 60.9 | Duo14 |
| ProteinMPNN$^D$ | -6.4 | 24.3 | 77.5 | -36.0 | 22.5 | 234.7 | Duo15 |
| | -17.7 | 42.6 | 44.2 | 63.4 | 87.8 | 141.3 | Duo16 |
| | 31.2 | 35.7 | 107.2 | -270.3 | 81.4 | 211.7 | Duo17 |
| | 40.4 | -14.2 | 93.0 | 27.4 | -297.5 | 148.0 | Duo18 |
| | 82.6 | 81.0 | 201.3 | 151.3 | -216.4 | 281.8 | Duo19 |
| | 87.3 | 47.6 | 218.2 | 104.0 | 221.3 | 621.8 | Duo20 |
| | 34.1 | 62.4 | 164.0 | -103.0 | -19.1 | 404.9 | Duo21 |
| | -28.5 | 6.4 | 76.3 | 251.4 | 251.0 | 604.1 | Duo22 |
| | -18.0 | 5.1 | 33.9 | -347.6 | -294.1 | 161.6 | Duo23 |
| | 45.1 | 46.0 | 50.7 | -215.8 | 87.5 | 167.5 | Duo24 |

* Mutant sequence pairs were scored on the WT template conformation.

**List S1:** Amino acid sequences of recombinant proteins and number of incorporated mutations compared with WT RMM.

| Protein | Mut nb | Amino acids sequence |
|---|---|---|
| RMM (WT) | | MSSNAIGLIETKGYVAALAAADAMVKAANVTITDRQQVGDGLVAVIVTGEVGAVKAATEAGAETASQVGELVSVHVIPRPHSELGAHFSVSSK |
| Duo1A | 20 | MSSNAIGAIQTKGTGAAIAAADAMVKAANVTLTSADTTGDGNVVVYVTGEVGAVKAATEAGAETASQDGELVAVYVTPRPHSELGAKRSVSSK |
| Duo1B | 18 | MSSNAIGIIITKGTVAADAAADAMVKAANVTDTIIDTTGDGNVLVLVTGEVGAVKAATEAGAETASQVGELLIVIVLPRPHSELGAVFSVSSK |
| Duo2A | 18 | MSSNAIGGIQTKGFAAALAAADAMVKAANVTVTAVVTTGDGEVKVYVTGEVGAVKAATEAGAETASQVGELLGVGVIPRPHSELGAIRSVSSK |
| Duo2B | 19 | MSSNAIGVITTKGFIAADAAADAMVKAANVTPTDLVTTGDGEVLVLVTGEVGAVKAATEAGAETASQLGELLTVVVLPRPHSELGAAFSVSSK |
| Duo3A | 26 | MSSNAKGAIQTKGWGAAIIAADAMIKAANVTLTSAKTTGGGNVAVYVTGEVGAVKAATEAGAETASQLGELVAVYVFPRPGSNHGAKRSVSSK |
| Duo3B | 20 | MSSNAIGIIITKGWVAADAAADAMEKAANVTDTDIKTTGGGNVLVLVTGEVGAVKAATEAGAETASQVGELLIVGVVPRPWSELGAVFSVSSK |
| Duo4A | 25 | MSSNAIGVIITKGFVAAVAAADAMVKAANVVLTSVYNTGDGQVLVLVTGEVGAVKAATEAGAETASQVGELLFVIVFPHPHEDLGAGADISSK |
| Duo4B | 24 | MSSNAIGLIATKGFGAALAAADAMVKAANVVGTPLYNTGDGQVVVFVTGEVGAVKAATEAGAETASQDGELVAVYVLPHPHEDLGAVLDISSK |
| Duo5A | 32 | MSSNAVGGIQTKGAGAAIGAADAMLKAANVVLTSAEVTGAGEVVVYVTGEVGAVKAATEAGAETASQGGELLAVAVIPHPLEIFGANRDISSK |
| Duo5B | 28 | MSSNALGIIITKGAVAADVAADAMGKAANVVGTSTEVTGAGEVLVLVTGEVGAVKAATEAGAETASQVGELLNVLVFPHPHEQLGAVFDISSK |
| Duo6A | 17 | MSSNAIGLISTKGFGAALAAADAMVKAANVTLTSGFNTGDGNVAVFVTGEVGAVKAATEAGAETASQAGELLAVHVLPRPHSELGAKLSVSSK |
| Duo6B | 18 | MSSNAIGVIVTKGFTAATAAADAMVKAANVTITSVFNTGDGNVLVLVTGEVGAVKAATEAGAETASQVGELYLVIVMPRPHSELGAIFSVSSK |
| Duo7A | 18 | MSSNAIGGIETKGAGAAIAAADAMVKAANVTLTDITNTGDGMVAVYVTGEVGAVKAATEAGAETASQAGELIAVAVFPRPHSELGATRSVSSK |
| Duo7B | 17 | MSSNAIGIITTKGAVAADAAADAMVKAANVTPTATTNTGDGMVLVLVTGEVGAVKAATEAGAETASQVGELINVIVIPRPHSELGAKFSVSSK |
| Duo8A | 23 | MSSNAVGGIETKGAGAAIAAADAMLKAANVTLTDITNTGGGMVAVYVTGEVGAVKAATEAGAETASQAGELIAVAVIPRPASIFGATRSVSSK |
| Duo8B | 19 | MSSNAIGIIVTKGAVAADVAADAMVKAANVTPTATTNTGGGMVGVLVTGEVGAVKAATEAGAETASQVGELINVIVIPRPHSELGAKFSVSSK |
| Duo9A | 19 | MSSNAIGLITTKGTGAALAAADAMVKAANVTVTSIKSSGDGNVTVFVTGEVGAVKAATEAGAETASQIGELLAVLVIPRPHSDLGAVLSVSSK |
| Duo9B | 22 | MSSNAIGVIVTKGTTAAVAAADAMVKAANVTLTSYKSSGDGNVLVTVTGEVGAVKAATEAGAETASQAGELIVVYVNPRPHSDLGAKASVSSK |
| Duo10A | 25 | MSSNAVGLITTKGIGAALDAADAMLKAANVTITSILSCGGGMCTVFVTGEVGAVKAATEAGAETASQIGELLAVLVLPRPSSTLGAVLSVSSK |
| Duo10B | 26 | MSSNAIGVIVTKGITAAVAAADAMTKAANVTLTSFLSCGGGMCLVTVTGEVGAVKAATEAGAETASQAGELLVVSVRPRPISQLGAKASVSSK |
| Duo11A | 19 | MSSNAIGIIITKGFVAAHAAADAMVKAANVTGTIIMTCGDGNVLVIVTGEVGAVKAATEAGAETASQVGELAYVGVLPRPHSDLGAVLSVSSK |
| Duo11B | 21 | MSSNAIGAIATKGFGAAIAAADAMVKAANVTLTAFMTCGDGNVVVYVTGEVGAVKAATEAGAETASQWGELVAVYVTPRPHSDLGAFDSVSSK |
| Duo12A | 22 | MSSNAVGLITTKGFGAALGAADAMVKAANVTITSIKNTGNGNVTVFVTGEVGAVKAATEAGAETASQAGELLAVAVLPRPHSDLGAVLSVSSK |
| Duo12B | 23 | MSSNAIGVIVTKGFTAAAVAADAMTKAANVTLTSAKNTGNGNVAVLVTGEVGAVKAATEAGAETASQVGELLIVLVFPRPHSQLGAKASVSSK |
| Duo13A | 18 | MSSNAIGLITTKGTGAALAAADAMVKAANVTVTVITKCGDGSVNVFVTGEVGAVKAATEAGAETASQIGELAAVLVIPRPHSELGAILSVSSK |
| Duo13B | 21 | MSSNAIGVIITKGTTAATAAADAMVKAANVTLTSITKCGDGSVLVLVTGEVGAVKAATEAGAETASQAGELIVVYVSPRPHSELGAAASVSSK |
| Duo14A | 20 | MSSNAIGGIQTKGGGAAIAAADAMVKAANVTVTGLRTTGDGEVVVYVTGEVGAVKAATEAGAETASQVGELLAVGVYPRPHSELGALRSVSSK |
| Duo14B | 19 | MSSNAIGIIITKGGVAADAAADAMVKAANVTGTSLRTTGDGEVLVLVTGEVGAVKAATEAGAETASQLGELLVVIVLPRPHSELGAAFSVSSK |

| | | |
|---|---|---|
| Duo15A | 17 | MSSNAIGIIETKGVTAATAAADAMVKAANVTQTDFRSDGDGSVLVLVTGEVGAVKAATEAGAETASQKGELLIVRVIPRPHSELGAAFSVSSK |
| Duo15B | 20 | MSSNAIGQIKTKGMAAAIAAADAMVKAANVTETAVRSDGDGSVAVFVTGEVGAVKAATEAGAETASQGGELQEVTVDPRPHSELGAKWSVSSK |
| Duo16A | 15 | MSSNAIGIIETKGDTAANAAADAMVKAANVTITGRQQSGDGQVTVLVTGEVGAVKAATEAGAETASQVGELMTVTVKPRPHSELGAVVSVSSK |
| Duo16B | 15 | MSSNAIGQIETKGWAAAIAAADAMVKAANVTITNLQQSGDGQVRVNVTGEVGAVKAATEAGAETASQVGELLQVQVVPRPHSELGAAFSVSSK |
| Duo17A | 22 | MSSNAVGIIETKGNVAADFAADAMLKAANVTITDVQRSGNGSDTVIVTGEVGAVKAATEAGAETASQRGELLTVTVPPRPWSAIGACFSVSSK |
| Duo17B | 23 | MSSNAVGRIETKGFAAAMMAADAMYKAANVTSTSVQRSGNGSVTVVVTGEVGAVKAATEAGAETASQVGELFCVGVEPRPHSKLGAKRSVSSK |
| Duo18A | 27 | MSSNAVGGIQTLGEGAATAAADAMVQAANVKTTDMKDNGNGHVTVIVTGEVGAVKAATEAGAETASQAGELMQVAVIPRPNMDLGAYFAASSK |
| Duo18B | 26 | MSSNAIGIIETKGFVAAMCAADAMVDAANVKLTAVKDNGNGHVLVRVTGEVGAVKAATEAGAETASQVGELREVLVIPRPLDYLGARQGISSK |
| Duo19A | 29 | MSSNAIGGIATMRNTAALKALNAMVAAANVTITSIDRDGDSGSTVWVTGEVGAVKAATEAGAETASQVGELRGVGVLPRPNEALGAFFKVSSK |
| Duo19B | 28 | MSSNAIGAITTSGAVAAVMAGDAMVTAANVTMTNWDRDGDSGVTVLVTGEVGAVKAATEAGAETASQMGELQEVFVEPRPTSTLGAAAWVSSK |
| Duo20A | 31 | MSSNAMGGIETLTFAAAIMAEAAMVAAANVVITAVLNQGDADVKVWVTGEVGAVKAATEAGAETASQVGELKRVDVEPRPDNDLGAVFDFSSK |
| Duo20B | 29 | MSSNAVGIIETKGEVAADKAAPAMVRAANVLFTAKLNQGDADTCVIVTGEVGAVKAATEAGAETASQVGELRTVTVAPRPISPIGAKVCISSK |
| Duo21A | 31 | MSSNARGGIQTYSVSAALAAATAMVKAANVTLTAVHDSGNAEHTVAVTGEVGAVKAATEAGAETASQVGELRQVAVNPRPHTLVGANIRLSSK |
| Duo21B | 30 | MSSNAIGSITTWGFVAAYEAEDAMVRAANVAPTALHDSGNAEQCVLVTGEVGAVKAATEAGAETASQVGELITVVVEPRPASSLGAAFDYSSK |
| Duo22A | 19 | MSSNAVGGIFTKGYSAALGAADAMCKAANVTITDRQQDGEGLVSVKVTGEVGAVKAATEAGAETASQVGELLQVGVNPRPDSRNGAVLSVSSK |
| Duo22B | 20 | MSSNAIGVIETKGDLAAEWAADAMLKAANVTPTSYQQDGEGLVTVLVTGEVGAVKAATEAGAETASQVGELLEVFVIPRPDSVIGARFSVSSK |
| Duo23A | 19 | MSSNAIGIIETKGIVAAIAAADAMLKAANVTITATRNDGDGRVLVRVTGEVGAVKAATEAGAETASQVGELLNVGVIPRPDSTLGALLSVSSK |
| Duo23B | 22 | MSSNAVGMIQTKGEGAAVVAADAMVKAANVTLTHVRNDGDGRVTVVVTGEVGAVKAATEAGAETASQWGELLQVHVIPRPHSDLGATWSVSSK |
| Duo24A | 22 | MSSNAIGLITTKGAVAAMFAADAMLKAANVTPTSRQSTGDGMDTVFVTGEVGAVKAATEAGAETASQIGELLEVAVNPRPNSSLGARWSVSSK |
| Duo24B | 23 | MSSNACGAIQTKGPTAAVMAADAMLKAANVTLTDVQSTGDGMVVVIVTGEVGAVKAATEAGAETASQVGELIEVGVLPRPNSKSGAIWSVSSK |

**List S2. BMC-H variant Duo DNA sequences for tripartite GFP assay.**

Fragment DNA sequences coding for A-GFP10 and B-GFP11 Duo monomers are provided with indication of flanking homology regions (blue lowercase letters) required for the Gibson assembly of each pair of monomers with the NdeI/SalI open vector-1. NdeI and SalI sites are colored purple and red, respectively. The complete fragment sequence is given for Duo1 monomers A and B, which are representative examples of GFP10 and GFP11-tagged organizations, respectively. Only the variable portion of the sequence, between NdeI and NotI sites (in blue), is given for all other Duo sequences.

Receptor Vector-1 derives from a pET26b-vector with insertion of the indicated DNA portion between BglII and BlpI sites (yellow boxes). This vector codes for WT RMM-GFP10/RMM-GFP11 and $His_6$-tagged GFP1-9, and serves as a positive interaction control. Coding regions are in bold letters, the GFP10 sequence in light green and GFP11 in dark green.

| Case | Sequence |
|---|---|
| Vector-1 (BglII/BlpI) | AGATCTCGATCCCGCGAAATTAATACGACTCACTATAGGGGAATTGTGAGCGGATAACAATTCCCCTCTAGAAATAagatttAAAtacttt<br>aagaaggagatatacatATGAGTAGTAACGCGATTGGTTTAATTGAAACGAAAGGATACGTCGCCGCACTGGCTGCTGCAGATGCTATG<br>GTAAAAGCTGCAAATGTGACCATCACCGACCGGCAGCAGGTTGGCGATGGCTTAGTGGCAGTGATCGTAACGGGTGAGGTTGGG<br>GCCGTAAAAGCTGCCACTGAAGCAGGCGCTGAAACTGCGTCGCAGGTTGGCGAGCTGGTTAGCGTGCATGTTATCCCACGTCCCC<br>ATTCGGAACTCGGCGCACATTTTAGCGTTAGCTCAAAAGGTGCGGCCGCATCAGAAGGAGGCGGTAGCGGGGGCCCTGGTTCGG<br>GAGGGGAAGGTTCTGCTGGGGGAGGGAGCGCTGGCGGGGGGTCTGATTTACCAGACGATCATTACCTGAGCACACAAACGATCC<br>TTTCGAAAGACCTGAACGCAAGCTGAggatcaattgtttaaGAAGGAGATATACCATGGCAAGTAGTAACGCGATTGGTTTAATTGAAA<br>CGAAAGGATACGTCGCCGCACTGGCTGCTGCAGATGCTATGGTAAAAGCTGCAAATGTGACCATCACCGACCGGCAGCAGGTTGG<br>CGATGGCTTAGTGGCAGTGATCGTAACGGGTGAGGTTGGGGCCGTAAAAGCTGCCACTGAAGCAGGCGCTGAAACTGCGTCGCA<br>GGTTGGCGAGCTGGTTAGCGTGCATGTTATCCCACGTCCCCATTCGGAACTCGGCGCACATTTTAGCGTTAGCTCAAAAGGATCCG<br>CAGGCAGCGGTGGAAGTCCGGGTGGCGGTTCAGGCGGTAGCGGCAGCTCTGCGAGCGGCGGCAGCACCAGCGAAAAACGCGAT<br>CACATGGTGCTGCTGGAATATGTGACCGCGGCGGGCATTACCGATGCGAGCTAATGACAAGTATGtcgactcctaggaaagctttCTCGA<br>GTTAACTCGTGAGCAATAACTAGCATAACCCCTTGGGGCCTCTAAACGGGTCTTGAGGGGTTTTTTGCTGAAAGTACACGGCCGCAT<br>AATCGAAATTAATACGACTCACTATAGGGGAATTGTGAGCGGATAACAATTCCCCTCTAGAATTAATTAAGTTTAACTTTAAGAAGGA<br>GATATACCTATGCGCAAAGGCGAAGAACTGTTTACCGGCGTGGTGCCGATTCTGATTGAACTGGATGGCGATGTGAACGGCCATA<br>AATTTTTTGTGCGCGGCGAAGGCGAAGGCGATGCGACCATTGGCAAACTGAGCCTGAAATTTATTTGCACCACCGGCAAACTGCC<br>GGTGCCGTGGCCGACCCTGGTGACCACCCTGACCTATGGCGTGCAGTGCTTTAGCCGCTATCCGGATCACATGAAACGCCATGATT<br>TTTTTAAAAGCGCGATGCCGGAAGGCTATGTGCAGGAACGCACCATTTATTTTAAAGATGATGGCACCTATAAAACCCGCGCGGA<br>AGTGAAATTTGAAGGCGATACCCTGGTGAACCGCATTGAACTGAAAGGCATTGATTTTAAAGAAGATGGCAACATTCTGGGCCAT<br>AAACTGGAATATAACTTTAACAGCCATAAAGTGTATATTACCGCGGATAAACAGAACAACGGCATTAAAGCGAACTTTACCATTCG<br>CCATAACGTGGAAGATGGCAGCGTGCAGCTGGCGGATCATTATCAGCAGAACACCCCGATTGGCGATGGCCCGGTGCTGCTGCCG<br>GATAACGGCAGCTCTGGTGCACATCACCATCACCATCATTAAGCGGCAGCACTTGTTACCGGTCACCTCTCGAGAAAACGCGTCGAG<br>AGCTGAG |
| **Sequence of fragments coding for A monomers for Gibson assembly** | |
| Duo1A | agatttAAAtactttaagaaggagatatacatATGAGTAGTAACGCGATTGGTGCTATTCAGACGAAAGGAACCGGGGCCGCAATCGCTGCT<br>GCAGATGCTATGGTAAAAGCTGCAAATGTGACCCTGACCAGCGCTGATACCACAGGCGATGGCAATGTGGTCGTGTACGTAACGG<br>GTGAGGTTGGGGCCGTAAAAGCTGCCACTGAAGCAGGCGCTGAAACTGCGTCGCAGGACGGCGAGCTGGTTGCGGTGTATGTTA<br>CCCCACGTCCCCATTCGGAACTCGGCGCAAAACGTAGCGTTAGCTCAAAAGGTGCGGCCGCATCAGAAGGAGGCGGTAGCGGGG<br>GCCCTGGTTCGGGAGGGGAAGGTTCTGCTGGGGGAGGGAGCGCTGGCGGGGGGTCTGATTTACCAGACGATCATTACCTGAGCA<br>CACAAACGATCCTTTCGAAAGACCTGAACGCAAGCTGATAAggatcaattgtttaa |
| **Only given portion between NdeI and NotI for remaining sequences** | |
| Duo2A | catATGAGTAGTAACGCGATTGGTGGGATTCAGACGAAAGGATTCGCTGCCGCACTGGCTGCTGCAGATGCTATGGTAAAAGCTGC<br>AAATGTGACCGTGACCGCGGTAGTGACCACAGGCGATGGCGAGGTGAAGGTGTACGTAACGGGTGAGGTTGGGGCCGTAAAAG<br>CTGCCACTGAAGCAGGCGCTGAAACTGCGTCGCAGGTTGGCGAGCTGTTAGGTGTGGGCGTTATCCCACGTCCCCATTCGGAACTC<br>GGCGCAATTCGTAGCGTTAGCTCAAAAGGTGCGGCCGC |
| Duo3A | catATGAGTAGTAACGCGAAGGGTGCTATTCAGACGAAAGGATGGGGGGCCGCAATTATCGCTGCAGATGCTATGATCAAAGCTG<br>CAAATGTGACCCTGACCAGCGCTAAAACCACAGGCGGCGGCAATGTGGCAGTGTACGTAACGGGTGAGGTTGGGGCCGTAAAAG<br>CTGCCACTGAAGCAGGCGCTGAAACTGCGTCGCAGTTAGGCGAGCTGGTTGCGGTGTATGTTTTCCCACGTCCCGGCTCGAACCAC<br>GGCGCAAAACGTAGCGTTAGCTCAAAAGGTGCGGCCGC |
| Duo4A | catATGAGTAGTAACGCGATTGGTGTAATTATTACGAAAGGATTCGTCGCCGCAGTGGCTGCTGCAGATGCTATGGTAAAAGCTGC<br>AAATGTGGTGCTGACCAGCGTATATAACACAGGCGATGGCCAAGTGTTGGTGCTGGTAACGGGTGAGGTTGGGGCCGTAAAAGC<br>TGCCACTGAAGCAGGCGCTGAAACTGCGTCGCAGGTTGGCGAGCTGTTATTCGTGATTGTTTTCCCACACCCCCATGAGGATCTCG<br>GCGCAGGCGCGGACATCAGCTCAAAAGGTGCGGCCGC |
| Duo5A | catATGAGTAGTAACGCGGTGGGTGGGATTCAGACGAAAGGAGCGGGGGCCGCAATCGGCGCTGCAGATGCTATGTTGAAAGCT<br>GCAAATGTGGTGCTGACCAGCGCTGAAGTGACAGGCGCGGGCGAGGTGGTCGTGTACGTAACGGGTGAGGTTGGGGCCGTAAA<br>AGCTGCCACTGAAGCAGGCGCTGAAACTGCGTCGCAGGGGGGCGAGCTGTTAGCGGTGGCGGTTATCCCACACCCCCTGGAGATT<br>TTCGGCGCAAACCGTGACATCAGCTCAAAAGGTGCGGCCGC |

| | |
|---|---|
| Duo6A | catATGAGTAGTAACGCGATTGGTTTAATTAGCACGAAAGGATTCGGGGCCGCACTGGCTGCTGCAGATGCTATGGTAAAAGCTGC<br>AAATGTGACCCTGACCAGCGGATTTAACACAGGCGATGGCAATGTGGCAGTGTTCGTAACGGGTGAGGTTGGGGCCGTAAAAGC<br>TGCCACTGAAGCAGGCGCTGAAACTGCGTCGCAGGCCGGCGAGCTGTTAGCGGTGCATGTTCTGCCACGTCCCCATTCGGAACTCG<br>GCGCAAAACTGAGCGTTAGCTCAAAAGGTGCGGCCGC |
| Duo7A | catATGAGTAGTAACGCGATTGGTGGGATTGAAACGAAAGGAGCGGGGGCCGCAATCGCTGCTGCAGATGCTATGGTAAAAGCT<br>GCAAATGTGACCCTGACCGACATAACCAACACAGGCGATGGCATGGTGGCAGTGTACGTAACGGGTGAGGTTGGGGCCGTAAAA<br>GCTGCCACTGAAGCAGGCGCTGAAACTGCGTCGCAGGCCGGCGAGCTGATCGCGGTGGCGGTTTTCCCACGTCCCCATTCGGAAC<br>TCGGCGCAACCCGTAGCGTTAGCTCAAAAGGTGCGGCCGC |
| Duo8A | catATGAGTAGTAACGCGGTGGGTGGGATTGAAACGAAAGGAGCGGGGGCCGCAATCGCTGCTGCAGATGCTATGTTGAAAGCT<br>GCAAATGTGACCCTGACCGACATAACCAACACAGGCGGCGGCATGGTGGCAGTGTACGTAACGGGTGAGGTTGGGGCCGTAAAA<br>GCTGCCACTGAAGCAGGCGCTGAAACTGCGTCGCAGGCCGGCGAGCTGATCGCGGTGGCGGTTATCCCACGTCCCGCGTCGATTT<br>TCGGCGCAACCCGTAGCGTTAGCTCAAAAGGTGCGGCCGC |
| Duo9A | catATGAGTAGTAACGCGATTGGTTTAATTACCACGAAAGGAACCGGGGCCGCACTGGCTGCTGCAGATGCTATGGTAAAAGCTGC<br>AAATGTGACCGTGACCAGCATAAAAAGCAGCGGCGATGGCAATGTGACGGTGTTCGTAACGGGTGAGGTTGGGGCCGTAAAAGC<br>TGCCACTGAAGCAGGCGCTGAAACTGCGTCGCAGATCGGCGAGCTGTTAGCGGTGCTGGTTATCCCACGTCCCCATTCGGATCTCG<br>GCGCAGTTCTGAGCGTTAGCTCAAAAGGTGCGGCCGC |
| Duo10A | catATGAGTAGTAACGCGGTGGGTTTAATTACCACGAAAGGAATTGGGGCCGCACTGGACGCTGCAGATGCTATGTTGAAAGCTG<br>CAAATGTGACCATCACCAGCATACTGAGCTGTGGCGGCGGCATGTGTACGGTGTTCGTAACGGGTGAGGTTGGGGCCGTAAAAG<br>CTGCCACTGAAGCAGGCGCTGAAACTGCGTCGCAGATCGGCGAGCTGTTAGCGGTGCTGGTTCTGCCACGTCCCAGCTCGACCCTC<br>GGCGCAGTTCTGAGCGTTAGCTCAAAAGGTGCGGCCGC |
| Duo11A | catATGAGTAGTAACGCGATTGGTATCATTATTACGAAAGGATTCGTCGCCGCACATGCTGCTGCAGATGCTATGGTAAAAGCTGC<br>AAATGTGACCGGCACCATCATAATGACCTGTGGCGATGGCAATGTGTTGGTGATCGTAACGGGTGAGGTTGGGGCCGTAAAAGC<br>TGCCACTGAAGCAGGCGCTGAAACTGCGTCGCAGGTTGGCGAGCTGGCCTACGTGGGCGTTCTGCCACGTCCCCATTCGGATCTCG<br>GCGCAGTTCTGAGCGTTAGCTCAAAAGGTGCGGCCGC |
| Duo12A | catATGAGTAGTAACGCGGTGGGTTTAATTACCACGAAAGGATTCGGGGCCGCACTGGGCGCTGCAGATGCTATGGTAAAAGCTG<br>CAAATGTGACCATCACCAGCATAAAAAACACAGGCAACGGCAATGTGACGGTGTTCGTAACGGGTGAGGTTGGGGCCGTAAAAG<br>CTGCCACTGAAGCAGGCGCTGAAACTGCGTCGCAGGCCGGCGAGCTGTTAGCGGTGGCGGTTCTGCCACGTCCCCATTCGGATCTC<br>GGCGCAGTTCTGAGCGTTAGCTCAAAAGGTGCGGCCGC |
| Duo13A | catATGAGTAGTAACGCGATTGGTTTAATTACCACGAAAGGAACCGGGGCCGCACTGGCTGCTGCAGATGCTATGGTAAAAGCTGC<br>AAATGTGACCGTGACCGTGATAACCAAATGTGGCGATGGCTCAGTGAATGTGTTCGTAACGGGTGAGGTTGGGGCCGTAAAAGC<br>TGCCACTGAAGCAGGCGCTGAAACTGCGTCGCAGATCGGCGAGCTGGCCGCGGTGCTGGTTATCCCACGTCCCCATTCGGAACTCG<br>GCGCAATTCTGAGCGTTAGCTCAAAAGGTGCGGCCGC |
| Duo14A | catATGAGTAGTAACGCGATTGGTGGGATTCAGACGAAAGGAGGCGGGGCCGCAATCGCTGCTGCAGATGCTATGGTAAAAGCTG<br>CAAATGTGACCGTGACCGGCCTTCGTACCACAGGCGATGGCGAGGTGGTCGTGTACGTAACGGGTGAGGTTGGGGCCGTAAAAG<br>CTGCCACTGAAGCAGGCGCTGAAACTGCGTCGCAGGTTGGCGAGCTGTTAGCGGTGGGCGTTTACCCACGTCCCCATTCGGAACTC<br>GGCGCACTGCGTAGCGTTAGCTCAAAAGGTGCGGCCGC |
| Duo15A | catATGAGTAGTAACGCGATTGGTATCATTGAAACGAAAGGAGTGACTGCCGCAACCGCTGCTGCAGATGCTATGGTAAAAGCTG<br>CAAATGTGACCCAAACCGACTTCCGTAGCGACGGCGATGGCTCAGTGTTGGTGCTGGTAACGGGTGAGGTTGGGGCCGTAAAAG<br>CTGCCACTGAAGCAGGCGCTGAAACTGCGTCGCAGAAGGGCGAGCTGTTAATCGTGCGTGTTATCCCACGTCCCCATTCGGAACTC<br>GGCGCAGCGTTTAGCGTTAGCTCAAAAGGTGCGGCCGC |
| Duo16A | catATGAGTAGTAACGCGATTGGTATCATTGAAACGAAAGGAGACACTGCCGCAAACGCTGCTGCAGATGCTATGGTAAAAGCTG<br>CAAATGTGACCATCACCGGCCGGCAGCAGAGCGGCGATGGCCAAGTGACGGTGCTGGTAACGGGTGAGGTTGGGGCCGTAAAA<br>GCTGCCACTGAAGCAGGCGCTGAAACTGCGTCGCAGGTTGGCGAGCTGATGACGGTGACCGTTAAACCACGTCCCCATTCGGAAC<br>TCGGCGCAGTGGTGAGCGTTAGCTCAAAAGGTGCGGCCGC |
| Duo17A | catATGAGTAGTAACGCGGTGGGTATCATTGAAACGAAAGGAAACGTCGCCGCAGATTTCGCTGCAGATGCTATGTTGAAAGCTG<br>CAAATGTGACCATCACCGACGTACAGCGTAGCGGCAACGGCTCAGACACGGTGATCGTAACGGGTGAGGTTGGGGCCGTAAAAG<br>CTGCCACTGAAGCAGGCGCTGAAACTGCGTCGCAGCGCGGCGAGCTGTTAACGGTGACCGTTCCGCCACGTCCCTGGTCGGCGAT<br>AGGCGCATGCTTTAGCGTTAGCTCAAAAGGTGCGGCCGC |
| Duo18A | catATGAGTAGTAACGCGGTGGGTGGGATTCAGACGCTGGGAGAGGGGGCCGCAACCGCTGCTGCAGATGCTATGGTACAGGCT<br>GCAAATGTGAAGACCACCGACATGAAAGATAATGGCAACGGCCACGTGACGGTGATCGTAACGGGTGAGGTTGGGGCCGTAAA<br>AGCTGCCACTGAAGCAGGCGCTGAAACTGCGTCGCAGGCCGGCGAGCTGATGCAAGTGGCGGTTATCCCACGTCCCAACATGGAT<br>CTCGGCGCATATTTTGCCGCGAGCTCAAAAGGTGCGGCCGC |
| Duo19A | catATGAGTAGTAACGCGATTGGTGGGATTGCGACGATGCGGAACACTGCCGCACTGAAGGCTTTAAACGCTATGGTAGCGGCTG<br>CAAATGTGACCATCACCAGCATAGATCGTGACGGCGATAGCGGGAGCACGGTGTGGGTAACGGGTGAGGTTGGGGCCGTAAAA<br>GCTGCCACTGAAGCAGGCGCTGAAACTGCGTCGCAGGTTGGCGAGCTGCGCGGTGTGGGCGTTCTGCCACGTCCCAACGAGGCGC<br>TCGGCGCATTTTTTAAGGTTAGCTCAAAAGGTGCGGCCGC |
| Duo20A | catATGAGTAGTAACGCGATGGGTGGGATTGAAACGCTGACTTTCGCTGCCGCAATTATGGCTGAGGCGGCTATGGTAGCGGCTGC<br>AAATGTGGTGATCACCGCGGTACTGAACCAAGGCGATGCGGACGTGAAGGTGTGGGTAACGGGTGAGGTTGGGGCCGTAAAAG<br>CTGCCACTGAAGCAGGCGCTGAAACTGCGTCGCAGGTTGGCGAGCTGAAGCGCGTGGATGTTGAGCCACGTCCCGATAATGATCT<br>CGGCGCAGTGTTTGACTTCAGCTCAAAAGGTGCGGCCGC |

| | |
|---|---|
| Duo21-A | catATGAGTAGTAACGCGCGTGGTGGGATTCAGACGTATAGTGTGAGCGCCGCACTGGCTGCTGCAACCGCTATGGTAAAAGCTGC<br>AAATGTGACCCTGACCGCGGTACATGATAGCGGCAACGCGGAGCACACGGTGGCGGTAACGGGTGAGGTTGGGGCCGTAAAAG<br>CTGCCACTGAAGCAGGCGCTGAAACTGCGTCGCAGGTTGGCGAGCTGCGCCAAGTGGCGGTTAACCCACGTCCCCATACTCTGGT<br>AGGCGCAAACATTCGCCTGAGCTCAAAAGGTGCGGCCGC |
| Duo22A | catATGAGTAGTAACGCGGTGGGTGGGATTTTTACGAAAGGATACAGCGCCGCACTGGGCGCTGCAGATGCTATGTGTAAAGCTG<br>CAAATGTGACCATCACCGACCGGCAGCAGGACGGCGAAGGCTTAGTGTCTGTGAAAGTAACGGGTGAGGTTGGGGCCGTAAAAG<br>CTGCCACTGAAGCAGGCGCTGAAACTGCGTCGCAGGTTGGCGAGCTGTTACAAGTGGGCGTTAACCCACGTCCCGATTCGCGTAA<br>TGGCGCAGTTCTGAGCGTTAGCTCAAAAGGTGCGGCCGC |
| Duo23A | catATGAGTAGTAACGCGATTGGTATCATTGAAACGAAAGGAATTGTCGCCGCAATCGCTGCTGCAGATGCTATGTTGAAAGCTGC<br>AAATGTGACCATCACCGCGACTCGTAACGACGGCGATGGCCGGGTGTTGGTGCGTGTAACGGGTGAGGTTGGGGCCGTAAAAGC<br>TGCCACTGAAGCAGGCGCTGAAACTGCGTCGCAGGTTGGCGAGCTGTTAAATGTGGGCGTTATCCCACGTCCCGATTCGACCCTCG<br>GCGCACTGCTGTCCGTTAGCTCAAAAGGTGCGGCCGC |
| Duo24A | catATGAGTAGTAACGCGATTGGTTTAATTACCACGAAAGGAGCGGTCGCCGCAATGTTCGCTGCAGATGCTATGTTGAAAGCTGC<br>AAATGTGACCCCGACCAGCCGGCAGAGCACAGGCGATGGCATGGACACGGTGTTCGTAACGGGTGAGGTTGGGGCCGTAAAAGC<br>TGCCACTGAAGCAGGCGCTGAAACTGCGTCGCAGATCGGCGAGCTGTTAGAGGTGGCGGTTAACCCACGTCCCAACTCCAGCCTC<br>GGCGCACGTTGGAGCGTTAGCTCAAAAGGTGCGGCCGC |
| Sequence of fragments coding for B monomers for Gibson assembly | |
| Duo1B | taaggatcaattgtttaagaaggagatatacatATGAGTAGTAACGCGATTGGTATCATTATTACGAAAGGAACCGTCGCCGCAGATGCTGCT<br>GCAGATGCTATGGTAAAAGCTGCAAATGTGACCGACACCATCATAGATACCACAGGCGATGGCAATGTGTTGGTGCTGGTAACGG<br>GTGAGGTTGGGGCCGTAAAAGCTGCCACTGAAGCAGGCGCTGAAACTGCGTCGCAGGTTGGCGAGCTGTTAATCGTGATTGTTCT<br>GCCACGTCCCCATTCGGAACTCGGCGCAGTGTTTAGCGTTAGCTCAAAAGGTGCGGCCGCAGGCAGCGGTGGCAGCCCGGGCGGC<br>GGCAGCGGCGGCAGCGGCAGCAGCGCGAGCGGCGGCAGCACCAGCGAAAAACGCGATCACATGGTGCTGCTGGAATATGTGAC<br>CGCGGCGGGCATTACCGATGCGAGCTAATGACAAGTATGtcgactcctaggaaagcttt |
| Only given portion between NdeI and NotI for remaining sequences | |
| Duo2B | catATGAGTAGTAACGCGATTGGTGTAATTACCACGAAAGGATTCATCGCCGCAGATGCTGCTGCAGATGCTATGGTAAAAGCTGC<br>AAATGTGACCCCGACCGACCTTGTGACCACAGGCGATGGCGAGGTGTTGGTGCTGGTAACGGGTGAGGTTGGGGCCGTAAAAGC<br>TGCCACTGAAGCAGGCGCTGAAACTGCGTCGCAGTTAGGCGAGCTGTTAACGGTGGTGGTTCTGCCACGTCCCCATTCGGAACTCG<br>GCGCAGCGTTTAGCGTTAGCTCAAAAGGTGCGGCCGC |
| Duo3B | catATGAGTAGTAACGCGATTGGTATCATTATTACGAAAGGATGGGTCGCCGCAGATGCTGCTGCAGATGCTATGGAGAAAGCTG<br>CAAATGTGACCGACACCGACATAAAAACCACAGGCGGCGGCAATGTGTTGGTGCTGGTAACGGGTGAGGTTGGGGCCGTAAAAG<br>CTGCCACTGAAGCAGGCGCTGAAACTGCGTCGCAGGTTGGCGAGCTGTTAATCGTGGGCGTTGTGCCACGTCCCTGGTCGGAACT<br>CGGCGCAGTGTTTAGCGTTAGCTCAAAAGGTGCGGCCGC |
| Duo4B | catATGAGTAGTAACGCGATTGGTTTAATTGCGACGAAAGGATTCGGGGCCGCACTGGCTGCTGCAGATGCTATGGTAAAAGCTGC<br>AAATGTGGTGGGCACCCCGCTTTATAACACAGGCGATGGCCAAGTGGTCGTGTTCGTAACGGGTGAGGTTGGGGCCGTAAAAGCT<br>GCCACTGAAGCAGGCGCTGAAACTGCGTCGCAGGACGGCGAGCTGGTTGCGGTGTATGTTCTGCCACACCCCCATGAGGATCTCG<br>GCGCAGTGCTGGACATCAGCTCAAAAGGTGCGGCCGC |
| Duo5B | catATGAGTAGTAACGCGCTGGGTATCATTATTACGAAAGGAGCGGTCGCCGCAGATGTCGCTGCAGATGCTATGGGGAAAGCTG<br>CAAATGTGGTGGGCACCAGCACTGAAGTGACAGGCGCGGGCGAGGTGTTGGTGCTGGTAACGGGTGAGGTTGGGGCCGTAAAA<br>GCTGCCACTGAAGCAGGCGCTGAAACTGCGTCGCAGGTTGGCGAGCTGTTAAATGTGCTGGTTTTCCCACACCCCCATGAGCAGCT<br>CGGCGCAGTGTTTGACATCAGCTCAAAAGGTGCGGCCGC |
| Duo6B | catATGAGTAGTAACGCGATTGGTGTAATTGTGACGAAAGGATTCACTGCCGCAACCGCTGCTGCAGATGCTATGGTAAAAGCTGC<br>AAATGTGACCATCACCAGCGTATTTAACACAGGCGATGGCAATGTGTTGGTGCTGGTAACGGGTGAGGTTGGGGCCGTAAAAGCT<br>GCCACTGAAGCAGGCGCTGAAACTGCGTCGCAGGTTGGCGAGCTGTACTTAGTGATTGTTATGCCACGTCCCCATTCGGAACTCGG<br>CGCAATTTTTAGCGTTAGCTCAAAAGGTGCGGCCGC |
| Duo7B | catATGAGTAGTAACGCGATTGGTATCATTACCACGAAAGGAGCGGTCGCCGCAGATGCTGCTGCAGATGCTATGGTAAAAGCTGC<br>AAATGTGACCCCGACCGCGACTACCAACACAGGCGATGGCATGGTGTTGGTGCTGGTAACGGGTGAGGTTGGGGCCGTAAAAGC<br>TGCCACTGAAGCAGGCGCTGAAACTGCGTCGCAGGTTGGCGAGCTGATCAATGTGATTGTTATCCCACGTCCCCATTCGGAACTCG<br>GCGCAAAATTTAGCGTTAGCTCAAAAGGTGCGGCCGC |
| Duo8B | catATGAGTAGTAACGCGATTGGTATCATTGTGACGAAAGGAGCGGTCGCCGCAGATGTCGCTGCAGATGCTATGGTAAAAGCTG<br>CAAATGTGACCCCGACCGCGACTACCAACACAGGCGGCGGCATGGTGGGTGTGCTGGTAACGGGTGAGGTTGGGGCCGTAAAAG<br>CTGCCACTGAAGCAGGCGCTGAAACTGCGTCGCAGGTTGGCGAGCTGATCAATGTGATTGTTATCCCACGTCCCCATTCGGAACTC<br>GGCGCAAAATTTAGCGTTAGCTCAAAAGGTGCGGCCGC |
| Duo9B | catATGAGTAGTAACGCGATTGGTGTAATTGTGACGAAAGGAACCACTGCCGCAGTGGCTGCTGCAGATGCTATGGTAAAAGCTG<br>CAAATGTGACCCTGACCAGCTACAAAAGCAGCGGCGATGGCAATGTGTTGGTGACCGTAACGGGTGAGGTTGGGGCCGTAAAAG<br>CTGCCACTGAAGCAGGCGCTGAAACTGCGTCGCAGGCCGGCGAGCTGATCGTTGTGTATGTTAACCCACGTCCCCATTCGGATCTC<br>GGCGCAAAAGCGAGCGTTAGCTCAAAAGGTGCGGCCGC |
| Duo10B | catATGAGTAGTAACGCGATTGGTGTAATTGTGACGAAAGGAATTACTGCCGCAGTGGCTGCTGCAGATGCTATGACTAAAGCTGC<br>AAATGTGACCCTGACCAGCTTCCTGAGCTGTGGCGGCGGCATGTGTTTGGTGACCGTAACGGGTGAGGTTGGGGCCGTAAAAGCT<br>GCCACTGAAGCAGGCGCTGAAACTGCGTCGCAGGCCGGCGAGCTGTTAGTTGTGAGCGTTCGTCCACGTCCCATTTCGCAGCTCGG<br>CGCAAAAGCGAGCGTTAGCTCAAAAGGTGCGGCCGC |

| | |
|---|---|
| Duo11B | catATGAGTAGTAACGCGATTGGTGCTATTGCGACGAAAGGATTCGGGGCCGCAATCGCTGCTGCAGATGCTATGGTAAAAGCTGC<br>AAATGTGACCCTGACCGCGTTCATGACCTGTGGCGATGGCAATGTGGTCGTGTACGTAACGGGTGAGGTTGGGGCCGTAAAAGCT<br>GCCACTGAAGCAGGCGCTGAAACTGCGTCGCAGTGGGGCGAGCTGGTTGCGGTGTATGTTACCCCACGTCCCCATTCGGATCTCG<br>GCGCATTTGATAGCGTTAGCTCAAAAGGTGCGGCCGC |
| Duo12B | catATGAGTAGTAACGCGATTGGTGTAATTGTGACGAAAGGATTCACTGCCGCAGCGGTCGCTGCAGATGCTATGACTAAAGCTGC<br>AAATGTGACCCTGACCAGCGCTAAAAACACAGGCAACGGCAATGTGGCAGTGCTGGTAACGGGTGAGGTTGGGGCCGTAAAAGC<br>TGCCACTGAAGCAGGCGCTGAAACTGCGTCGCAGGTTGGCGAGCTGTTAATCGTGCTGGTTTTCCCACGTCCCCATTCGCAGCTCG<br>GCGCAAAAGCGAGCGTTAGCTCAAAAGGTGCGGCCGC |
| Duo13B | catATGAGTAGTAACGCGATTGGTGTAATTATTACGAAAGGAACCACTGCCGCAACCGCTGCTGCAGATGCTATGGTAAAAGCTGC<br>AAATGTGACCCTGACCAGCATAACCAAATGTGGCGATGGCTCAGTGTTGGTGCTGGTAACGGGTGAGGTTGGGGCCGTAAAAGC<br>TGCCACTGAAGCAGGCGCTGAAACTGCGTCGCAGGCCGGCGAGCTGATCGTTGTGTATGTTAGCCCACGTCCCCATTCGGAACTCG<br>GCGCAGCGGCGAGCGTTAGCTCAAAAGGTGCGGCCGC |
| Duo14B | catATGAGTAGTAACGCGATTGGTATCATTATTACGAAAGGAGGCGTCGCCGCAGATGCTGCTGCAGATGCTATGGTAAAAGCTGC<br>AAATGTGACCGGCACCAGCCTTCGTACCACAGGCGATGGCGAGGTGTTGGTGCTGGTAACGGGTGAGGTTGGGGCCGTAAAAGC<br>TGCCACTGAAGCAGGCGCTGAAACTGCGTCGCAGTTAGGCGAGCTGTTAGTTGTGATTGTTCTGCCACGTCCCCATTCGGAACTCG<br>GCGCAGCGTTTAGCGTTAGCTCAAAAGGTGCGGCCGC |
| Duo15B | catATGAGTAGTAACGCGATTGGTCAAATTAAAACGAAAGGAATGGCTGCCGCAATCGCTGCTGCAGATGCTATGGTAAAAGCTG<br>CAAATGTGACCGAGACCGCGGTACGTAGCGACGGCGATGGCTCAGTGGCAGTGTTCGTAACGGGTGAGGTTGGGGCCGTAAAAG<br>CTGCCACTGAAGCAGGCGCTGAAACTGCGTCGCAGGGGGGCGAGCTGCAAGAGGTGACCGTTGACCCACGTCCCCATTCGGAACT<br>CGGCGCAAAATGGAGCGTTAGCTCAAAAGGTGCGGCCGC |
| Duo16B | catATGAGTAGTAACGCGATTGGTCAAATTGAAACGAAAGGATGGGCTGCCGCAATCGCTGCTGCAGATGCTATGGTAAAAGCTG<br>CAAATGTGACCATCACCAATCTTCAGCAGAGCGGCGATGGCCAAGTGCGCGTGAATGTAACGGGTGAGGTTGGGGCCGTAAAAG<br>CTGCCACTGAAGCAGGCGCTGAAACTGCGTCGCAGGTTGGCGAGCTGTTACAAGTGCAGGTTGTGCCACGTCCCCATTCGGAACTC<br>GGCGCAGCGTTTAGCGTTAGCTCAAAAGGTGCGGCCGC |
| Duo17B | catATGAGTAGTAACGCGGTGGGTCGCATTGAAACGAAAGGATTCGCTGCCGCAATGATGGCTGCAGATGCTATGTACAAAGCTG<br>CAAATGTGACCAGCACCAGCGTACAGCGTAGCGGCAACGGCTCAGTGACGGTGGTGGTAACGGGTGAGGTTGGGGCCGTAAAA<br>GCTGCCACTGAAGCAGGCGCTGAAACTGCGTCGCAGGTTGGCGAGCTGTTCTGTGTGGGCGTTGAGCCACGTCCCCATTCGAAACT<br>CGGCGCAAAACGTAGCGTTAGCTCAAAAGGTGCGGCCGC |
| Duo18B | catATGAGTAGTAACGCGATTGGTATCATTGAAACGAAAGGATTCGTCGCCGCAATGTGTGCTGCAGATGCTATGGTAGATGCTGC<br>AAATGTGAAGCTGACCGCGGTAAAAGATAATGGCAACGGCCACGTGTTGGTGCGTGTAACGGGTGAGGTTGGGGCCGTAAAAGC<br>TGCCACTGAAGCAGGCGCTGAAACTGCGTCGCAGGTTGGCGAGCTGCGCGAGGTGCTGGTTATCCCACGTCCCCTGGACTATCTCG<br>GCGCACGTCAGGGTATCAGCTCAAAAGGTGCGGCCGC |
| Duo19B | catATGAGTAGTAACGCGATTGGTGCTATTACCACGAGCGGAGCGGTCGCCGCAGTGATGGCTGGCGATGCTATGGTAACCGCTGC<br>AAATGTGACCATGACCAACTGGGATCGTGACGGCGATAGCGGGGTGACGGTGCTGGTAACGGGTGAGGTTGGGGCCGTAAAAG<br>CTGCCACTGAAGCAGGCGCTGAAACTGCGTCGCAGATGGGCGAGCTGCAAGAGGTGTTTGTTGAGCCACGTCCCACCTCGACCCT<br>CGGCGCAGCGGCGTGGGTTAGCTCAAAAGGTGCGGCCGC |
| Duo20B | catATGAGTAGTAACGCGGTGGGTATCATTGAAACGAAAGGAGAGGTCGCCGCAGATAAGGCTGCACCGGCTATGGTACGTGCTG<br>CAAATGTGCTGTTCACCGCGAAGCTGAACCAAGGCGATGCGGACACATGTGTGATCGTAACGGGTGAGGTTGGGGCCGTAAAAG<br>CTGCCACTGAAGCAGGCGCTGAAACTGCGTCGCAGGTTGGCGAGCTGCGCACGGTGACCGTTGCGCCACGTCCCATTTCGCCGATA<br>GGCGCAAAAGTGTGTATCAGCTCAAAAGGTGCGGCCGC |
| Duo21B | catATGAGTAGTAACGCGATTGGTAGTATTACCACGTGGGGATTCGTCGCCGCATACGAGGCTGAGGATGCTATGGTACGTGCTGC<br>AAATGTGGCGCCGACCGCGCTTCATGATAGCGGCAACGCGGAGCAATGTGTGCTGGTAACGGGTGAGGTTGGGGCCGTAAAAGC<br>TGCCACTGAAGCAGGCGCTGAAACTGCGTCGCAGGTTGGCGAGCTGATCACGGTGGTGGTTGAGCCACGTCCCGCGTCGAGCCTC<br>GGCGCAGCGTTTGACTACAGCTCAAAAGGTGCGGCCGC |
| Duo22B | catATGAGTAGTAACGCGATTGGTGTAATTGAAACGAAAGGAGACTTGGCCGCAGAATGGGCTGCAGATGCTATGTTGAAAGCTG<br>CAAATGTGACCCCGACCAGCTACCAGCAGGACGGCGAAGGCTTAGTGACGGTGCTGGTAACGGGTGAGGTTGGGGCCGTAAAAG<br>CTGCCACTGAAGCAGGCGCTGAAACTGCGTCGCAGGTTGGCGAGCTGTTAGAGGTGTTTGTTATCCCACGTCCCGATTCGGTGATA<br>GGCGCACGTTTTAGCGTTAGCTCAAAAGGTGCGGCCGC |
| Duo23B | catATGAGTAGTAACGCGGTGGGTATGATTCAGACGAAAGGAGAGGGGGCCGCAGTGGTCGCTGCAGATGCTATGGTAAAAGCT<br>GCAAATGTGACCCTGACCCACGTACGTAACGACGGCGATGGCCGGGTGACGGTGGTGGTAACGGGTGAGGTTGGGGCCGTAAAA<br>GCTGCCACTGAAGCAGGCGCTGAAACTGCGTCGCAGTGGGGCGAGCTGTTACAAGTGCATGTTATCCCACGTCCCCATTCGGATCT<br>CGGCGCAACCTGGAGCGTTAGCTCAAAAGGTGCGGCCGC |
| Duo24B | catATGAGTAGTAACGCGTGTGGTGCTATTCAGACGAAAGGACCGACTGCCGCAGTGATGGCTGCAGATGCTATGTTGAAAGCTGC<br>AAATGTGACCCTGACCGACGTACAGAGCACAGGCGATGGCATGGTGGTCGTGATCGTAACGGGTGAGGTTGGGGCCGTAAAAGC<br>TGCCACTGAAGCAGGCGCTGAAACTGCGTCGCAGGTTGGCGAGCTGATCGAGGTGGGCGTTCTGCCACGTCCCAACTCGAAATCA<br>GGCGCAATTTGGAGCGTTAGCTCAAAAGGTGCGGCCGC |

**List S3. Duo DNA sequences for Flag/His$_6$ assay.**

Next DNA sequences were ordered directly cloned between BglII and XhoI sites of the pET24(+) vector (Kanamycin resistance) from Twist Biosciences. Flag-tag is colored in dark blue, other details are like for List S2.

| Case | Sequence |
|---|---|
| Duo2A/B | taatacgactcactataggggatctagatccgtatgacataaggaggtgaacatATGTCGTCGAACGCAATCGGTGGCATTCAGACTAAAGGTTTCGC<br>TGCGGCACTCGCCGCAGCTGACGCCATGGTTAAAGCAGCCAATGTCACTGTGACTGCTGTCGTAACGACGGGTGACGGTGAGGT<br>AAAGGTTTACGTTACCGGAGAGGTGGGCGCAGTTAAAGCTGCAACAGAGGCGGGAGCGGAAACGGCGAGCCAAGTTGGGGAG<br>CTATTGGGCGTAGGAGTCATACCGCGGCCGCACTCGGAGTTGGGTGCCATAAGATCCGTTAGCTCAAAGGGGAGCGGAAGTGG<br>GGCCGATTACAAAGATGACGATGACAAGTAAgtctgccaccctaagggggtcattgaATGAGCTCGAACGCTATCGGAGTAATAACGACA<br>AAGGGCTTCATCGCGGCTGACGCTGCGGCGGATGCCATGGTGAAGGCTGCTAACGTGACTCCCACAGACTTAGTGACCACGGGT<br>GATGGTGAAGTCCTAGTCTTGGTAACCGGAGAGGTCGGCGCCGTCAAGGCAGCAACGGAGGCAGGAGCCGAGACGGCCTCCCA<br>ATTAGGGGAGTTGCTGACTGTCGTTGTACTTCCCCGGCCGCATTCCGAACTCGGTGCAGCGTTCTCCGTTAGCTCAAAAGGTAGC<br>GGAAGTGGTCATCACCACCACCATCACTAAtaacaaagctagcataaccccttggggcctctaaacgggtcttgaggggttttttg |
| Duo3A/B | taatacgactcactataggggatctagagccccgaaccttagggaggtaaacatATGAGTTCTAATGCAAAGGGCGCCATTCAGACCAAAGGCTGGG<br>GCGCGGCAATCATCGCGGCGGACGCCATGATTAAAGCTGCTAACGTCACATTAACATCCGCGAAGACAACTGGGGGCGGGAAT<br>GTTGCAGTGTACGTCACTGGCGAAGTGGGTGCGGTTAAAGCGGCCACAGAAGCGGGAGCAGAAACGGCAAGTCAGCTTGGAG<br>AGCTTGTTGCAGTCTATGTGTTTCCCCGGCCCGGGAGTAACCACGGCGCTAAACGCTCAGTGAGCAGCAAGGGGTCAGGGTCCG<br>GGGCCGATTACAAAGACGACGACGACAAATGAaatagacgggctaaggaggttcgacgATGAGTTCTAACGCGATCGGGATCATCATTA<br>CTAAAGGGTGGGTAGCAGCTGATGCTGCGGCGGACGCCATGGAAAAGGCGGCAAACGTTACTGACACGGACATAAAGACGAC<br>AGGCGGCGGGAACGTTTTAGTTTTAGTTACCGGAGAGGTGGGAGCAGTAAAGGCCGCTACAGAGGCCGGTGCTGAAACCGCGT<br>CGCAGGTCGGTGAATTATTAATTGTCGGTGTCGTTCCGCGGCCGTGGTCAGAGCTGGGCGCTGTATTCAGCGTATCTTCTAAGGG<br>CTCAGGTAGTGGGCATCATCACCACCATCACTAAtaacaaagctagcataaccccttggggcctctaaacgggtcttgaggggttttttg |
| Duo4A/B | taatacgactcactataggggatctagaggaaaaaccccctaaggagggacacatATGAGTTCTAATGCAATTGGAGTGATCATCACAAAAGGATTCG<br>TAGCTGCGGTGGCCGCCGCTGATGCTATGGTCAAAGCAGCGAACGTGGTTTTAACTTCCGTGTATAACACTGGGGACGGGCAGG<br>TCCTCGTATTAGTAACCGGTGAGGTCGGTGCAGTAAAGGCCGCGACGGAGGCGGGAGCAGAAACTGCGTCCCAAGTCGGTGAG<br>TTATTGTTCGTAATCGTCTTCCCACATCCACACGAAGATCTCGGGGCCGGCGCTGACATTAGTTCCAAAGGCTCCGGAAGTGGAG<br>CGGACTACAAAGACGATGATGATAAATGAcccgtcattgataaggaggtccaagtATGAGCTCGAATGCGATCGGATTGATCGCAACGA<br>AAGGTTTCGGTGCGGCCTTGGCAGCAGCAGACGCCATGGTGAAGGCGGCTAACGTCGTTGGTACACCGTTGTACAACACTGGGG<br>ACGGACAAGTAGTTGTCTTTGTTACAGGCGAAGTAGGGGCTGTAAAGGCTGCCACAGAAGCAGGTGCGGAGACTGCATCCCAA<br>GACGGTGAGCTCGTCGCCGTGTACGTGTTGCCTCACCCGCACGAGGACCTCGGAGCGGTACTGGACATTTCTTCGAAGGGGTCTG<br>GTAGCGGACACCATCATCACCATCATTAAtaacaaagctagcataaccccttggggcctctaaacgggtcttgaggggttttttg |
| Duo6A/B | taatacgactcactataggggatctagagagcccgggattaaggaggacggcatATGTCCAGCAACGCAATCGGGTTAATAAGCACGAAAGGTTTCG<br>GCGCCGCACTGGCTGCCGCTGATGCTATGGTTAAAGCTGCGAATGTGACCCTGACCAGCGGATTTAATACAGGTGATGGTAATG<br>TGGCAGTGTTCGTAACGGGCGAAGTTGGTGCCGTCAAAGCTGCAACTGAAGCCGGCGCTGAAACCGCGTCGCAGGCCGGCGAG<br>CTGTTAGCGGTGCATGTTCTGCCACGCCCCCACTCTGAATTGGGCGCAAAACTGAGCGTGTCTAGTAAAGGATCGGGCAGCGGG<br>GCCGATTATAAGGATGACGATGACAAGTGAggttcgctgaataaggaggtcagggcATGTCGAGCAACGCCATTGGGGTCATTGTCACTA<br>AAGGCTTTACTGCCGCAACCGCCGCTGCAGATGCTATGGTAAAAGCTGCAAATGTGACCATCACCAGCGTATTTAACACAGGCG<br>ATGGCAATGTGTTGGTGCTGGTAACCGGTGAGGTGGGGGCGGTAAAGGCTGCCACGGAAGCAGGCGCAGAAACTGCGAGTCA<br>AGTTGGCGAGCTGTACTTAGTGATTGTTATGCCGCGTCCTCATTCGGAACTCGGTGCAATTTTTAGCGTTAGCTCAAAGGGAAGC<br>GGTAGCGGTCATCATCATCACCATCATTAAtaacaaagctagcataaccccttggggcctctaaacgggtcttgaggggttttttg |
| Duo7A/B | taatacgactcactataggggatctagatccgtatgacataaggaggtgaacatATGTCGTCGAACGCAATCGGTGGCATTGAGACTAAAGGTGCGG<br>GGGCCGCAATCGCTGCCGCTGATGCTATGGTTAAAGCTGCGAATGTGACCCTGACCGACATAACCAATACAGGTGATGGTATGG<br>TGGCAGTGTACGTAACGGGCGAAGTTGGTGCCGTCAAAGCTGCAACTGAAGCCGGCGCTGAAACCGCGTCGCAGGCCGGCGAG<br>CTGATCGCGGTGGCGGTTTTCCCACGCCCCCACTCTGAATTGGGCGCAACCCGTAGCGTGAGCTCAAAAGGCTCCGGAAGTGGA<br>GCGGACTACAAAGACGATGATGATAAATGAcccgtcattgataaggaggtccaagtATGAGCTCGAATGCGATCGGAATCATCACAACG<br>AAAGGTGCGGTCGCCGCAGACGCCGCTGCAGATGCTATGGTAAAAGCTGCAAATGTGACCCCGACCGCGACTACCAACACAGGC<br>GATGGCATGGTGTTGGTGCTGGTAACCGGTGAGGTGGGGGCGGTAAAGGCTGCCACGGAAGCAGGCGCAGAAACTGCGAGTC<br>AAGTTGGCGAGCTGATCAATGTGATTGTTATCCCGCGTCCTCATTCGGAACTCGGTGCAAAATTTAGCGTTAGCTCAAAGGGGTC<br>TGGTAGCGGACACCATCATCACCATCATTAAtaacaaagctagcataaccccttggggcctctaaacgggtcttgaggggttttttg |
| Duo9A/B | taatacgactcactataggggatctagagagcccgggattaaggaggacggcatATGTCCAGCAACGCAATCGGGTTAATAACCACGAAAGGTACA<br>GGCGCCGCGCTTGCAGCCGCAGATGCGATGGTCAAAGCGGCTAACGTGACAGTAACGAGCATTAAGTCCTCAGGGGATGGTAA<br>TGTGACAGTCTTCGTAACGGGTGAAGTTGGGGCCGTTAAGGCTGCAACTGAGGCAGGAGCTGAGACTGCCTCCCAGATTGGAG<br>AGTTGCTTGCTGTCTTGGTTATCCCTAGACCGCACTCAGACCTGGGCGCAGTTCTAAGTGTCTCTAGTAAAGGATCGGGCAGCGG<br>GGCCGATTATAAGGATGACGATGACAAGTGAggttcgctgaataaggaggtcagggcATGTCGAGCAACGCCATTGGGGTCATTGTCACT<br>AAAGGCACCACAGCTGCCGTCGCTGCCGCGGATGCCATGGTCAAGGCAGCCAATGTGACCTTGACTTCCTACAAGTCTAGCGGA<br>GACGGAAACGTCTTAGTGACAGTGACCGGAGAAGTGGGAGCCGTGAAAGCAGCCACTGAGGCTGGTGCCGAGACCGCAAGCC<br>AAGCCGGTGAGCTGATCGTAGTCTACGTGAATCCAAGACCTCATAGTGATCTAGGCGCCAAAGCTTCAGTGTCTAGCAAGGGAA<br>GCGGTAGCGGTCATCATCATCACCATCATTAAtaacaaagctagcataaccccttggggcctctaaacgggtcttgaggggttttttg |
| Duo10A/B | taatacgactcactataggggatctagagacgagggagataaggaggcgtgcatATGAGCAGCAACGCTGTGGGGTTGATAACTACCAAGGGAATA<br>GGAGCGGCATTAGACGCAGCTGACGCCATGCTGAAGGCCGCGAACGTTACCATTACGTCTATTCTCAGCTGCGGTGGTGGTATG<br>TGTACGGTGTTCGTTACCGGAGAAGTCGGAGCGGTAAAGGCCGCTACAGAAGCAGGAGCTGAAACTGCTAGTCAGATTGGAGA<br>GCTGTTAGCTGTTTTAGTTTTACCTCGTCCCTCCAGCACCCTCGGAGCCGTTCTAAGCGTGAGCTCCAAGGGGAGTGGATCCGGCG<br>CAGACTATAAAGATGATGATGATAAGTAAgaaaagaagtaaaaggaggcagatacATGAGCTCAAACGCTATAGGCGTAATCGTAACA<br>AAAGGAATCACGGCGGCCGTCGCAGCGGCGGATGCCATGACCAAGGCGGCCAACGTTACCTTGACTAGCTTCTTAAGCTGCGGC<br>GGTGGAATGTGTCTGGTGACGGTTACAGGTGAGGTTGGGGCCGTCAAAGCGGCTACCGAGGCCGGTGCCGAGACAGCGTCACA<br>GGCTGGAGAGCTTCTAGTCGTCAGCGTAAGACCGCGTCCTATCAGTCAGCTTGGGGCCAAAGCTAGTGTGTCTAGCAAAGGTAG<br>TGGTTCGGGCCACCATCATCACCATCATTGAtaacaaagctagcataaccccttggggcctctaaacgggtcttgaggggttttttg |

| | |
|---|---|
| Duo11A/B | taatacgactcactataggggatctagagaacgagtaatccaggaggtctgcatATGAGCTCTAATGCTATTGGGATCATAATCACCAAGGGCTTTGT<br>CGCCGCACATGCCGCAGCTGATGCAATGGTGAAGGCGGCGAATGTCACGGGCACGATCATTATGACCTGCGGTGACGGAAACG<br>TGTTAGTGATCGTTACCGGAGAAGTAGGCGCAGTGAAGGCCGCTACCGAGGCTGGCGCAGAGACTGCTTCGCAAGTCGGGGAG<br>CTAGCCTATGTGGGAGTTTTACCTCGGCCTCACTCAGACTTGGGCGCGGTACTATCAGTGTCATCAAAGGGGTCTGGTAGTGGCG<br>CCGACTACAAAGATGACGACGACAAGTAAagtcagagcggtaaggaggtccctttATGAGCAGTAACGCTATAGGTGCTATAGCTACAAA<br>AGGCTTTGGAGCAGCAATTGCGGCGGCTGACGCCATGGTCAAGGCGGCCAACGTAACGCTCACGGCGTTTATGACCTGCGGGG<br>ACGGAAATGTAGTTGTGTATGTGACTGGTGAGGTAGGTGCGGTAAAAGCCGCCACTGAGGCTGGGGCAGAAACAGCGTCCCAG<br>TGGGGAGAGCTCGTAGCGGTATACGTGACACCCCGCCCGCATTCAGATCTCGGTGCCTTCGACAGCGTCTCATCCAAGGGCAGTG<br>GCAGTGGGCATCACCACCACCACCATTAAtaacaaagctagcataaccccttggggcctctaaacgggtcttgaggggttttttg |
| Duo13A/B | taatacgactcactataggggatctagagagcccgggattaaggaggacggcatATGTCCAGCAACGCAATCGGGTTAATAACCACGAAAGGTACA<br>GGCGCCGCGCTTGCTGCCGCTGATGCTATGGTTAAAGCTGCGAATGTGACCGTGACCGTGATAACCAAGTGTGGTGATGGTTCA<br>GTGAATGTGTTCGTAACGGGCGAAGTTGGTGCCGTCAAAGCTGCAACTGAAGCCGGCGCTGAAACCGCGTCGCAGATCGGCGA<br>GCTGGCCGCGGTGCTGGTTATCCCACGCCCCCACTCTGAATTGGGCGCAATTCTGAGCGTGAGCTCAAAAGGATCGGGCAGCGG<br>GGCCGATTATAAGGATGACGATGACAAGTGAggttcgctgaataaggaggtcagggcATGTCGAGCAACGCCATTGGGGTCATTATTACT<br>AAAGGCACCACAGCTGCCACCGCTGCCGCAGATGCTATGGTAAAAGCTGCAAATGTGACCCTGACCAGCATAACCAAATGTGGC<br>GATGGCTCAGTGTTGGTGCTGGTAACCGGTGAGGTGGGGGCGGTAAAGGCTGCCACGGAAGCAGGCGCAGAAACTGCGAGTC<br>AAGCCGGCGAGCTGATCGTTGTGTATGTTAGCCCGCGTCCTCATTCGGAACTCGGTGCAGCGGCGAGCGTTAGCTCAAAGGGAA<br>GCGGTAGCGGTCATCATCATCACCATCATTAAtaacaaagctagcataaccccttggggcctctaaacgggtcttgaggggttttttg |
| Duo15A/B | taatacgactcactataggggatctagaggcccttttccaaaggaggttaacatATGAGTTCAAACGCAATCGGGATCATAGAGACAAAAGGAGTGA<br>CTGCGGCCACCGCTGCCGCTGATGCTATGGTTAAAGCTGCGAATGTGACCCAAACCGACTTCCGTAGCGACGGTGATGGTTCAGT<br>GTTGGTGCTGGTAACGGGCGAAGTTGGTGCCGTCAAAGCTGCAACTGAAGCCGGCGCTGAAACCGCGTCGCAGAAGGGCGAGC<br>TGTTAATCGTGCGTGTTATCCCACGCCCCCACTCTGAATTGGGCGCAGCGTTTAGCGTGAGCTCAAAAGGGTCAGGGTCCGGGGC<br>CGATTACAAAGACGACGACGACAAATGAaatagacgggctaaggaggttcgacgATGAGTTCTAACGCGATCGGGCAGATCAAAACTAA<br>AGGGATGGCTGCCGCAATCGCCGCTGCAGATGCTATGGTAAAAGCTGCAAATGTGACCGAGACCGCGGTACGTAGCGACGGCG<br>ATGGCTCAGTGGCAGTGTTCGTAACCGGTGAGGTGGGGGCGGTAAAGGCTGCCACGGAAGCAGGCGCAGAAACTGCGTCGCA<br>GGGGGGCGAGCTGCAAGAGGTGACCGTTGACCCGCGTCCACATTCGGAACTCGGCGCAAAATGGTCCGTGTCTTCAAAAGGTA<br>GCGGAAGTGGTCATCACCACCACCATCACTAAtaacaaagctagcataaccccttggggcctctaaacgggtcttgaggggttttttg |
| Duo23A/B | taatacgactcactataggggatctagaggcccttttccaaaggaggttaacatATGAGTTCAAACGCAATCGGGATCATAGAGACAAAAGGAATTG<br>TCGCGGCCATCGCGGCGGCGGACGCGATGTTAAAGGCGGCAAATGTGACTATTACTGCAACACGCAATGACGGTGATGGAAGA<br>GTGCTTGTTCGCGTCACAGGCGAGGTGGGTGCAGTTAAGGCAGCGACGGAGGCAGGCGCGGAGACTGCGTCGCAGGTCGGCG<br>AGTTGTTAAACGTCGGAGTGATTCCTAGGCCTGACAGCACCCTGGGTGCGCTGCTGAGCGTTAGCTCAAAAGGCAGCGGAAGCG<br>GTGCCGACTATAAGGACGACGATGACAAGTAActcgtttcacctaaggagggactcagATGAGCAGCAATGCGGTGGGAATGATCCAAA<br>CCAAAGGCGAGGGCGCTGCAGTAGTAGCCGCAGATGCTATGGTCAAGGCCGCCAACGTTACACTAACCCACGTCCGCAACGACG<br>GAGACGGTAGGGTGACGGTGGTTGTCACCGGAGAAGTTGGAGCCGTAAAAGCAGCCACGGAAGCAGGAGCGGAGACTGCGTC<br>TCAGTGGGGAGAGCTTCTCCAAGTTCATGTCATCCCCAGGCCTCACTCCGACTTGGGTGCGACATGGAGTGTCTCCAGTAAGGGG<br>TCTGGATCGGGCCATCACCATCATCACCATTAAtaacaaagctagcataaccccttggggcctctaaacgggtcttgaggggttttttg |
| Duo24A/B | taatacgactcactataggggatctagagagcccgggattaaggaggacggcatATGTCCAGCAACGCAATCGGGTTAATAACCACGAAAGGTGCA<br>GTCGCCGCAATGTTCGCCGCTGATGCAATGTTGAAAGCAGCGAATGTGACCCCGACCAGCCGGCAGAGCACAGGTGATGGTATG<br>GACACGGTGTTCGTAACGGGCGAAGTTGGTGCCGTCAAAGCTGCAACTGAAGCCGGCGCTGAAACCGCGTCGCAGATCGGCGA<br>GCTGTTAGAGGTGGCGGTTAACCCACGCCCCAACTCTAGCTTGGGCGCACGTTGGAGCGTGAGCAGCAAAGGCAGCGGAAGCG<br>GTGCCGACTATAAGGACGACGATGACAAGTAActcgtttcacctaaggagggactcagATGAGCAGCAATGCGTGTGGAGCTATTCAGAC<br>AAAAGGACCGACTGCCGCAGTCATGGCTGCAGATGCTATGTTAAAAGCTGCAAACGTGACCCTGACCGACGTACAGAGCACAG<br>GCGATGGCATGGTGGTCGTGATCGTAACCGGTGAGGTGGGGGCGGTAAAGGCTGCCACGGAAGCAGGCGCAGAAACTGCGAG<br>TCAAGTTGGCGAGCTGATCGAGGTGGGCGTTCTGCCGCGTCCTAACTCGAAATCAGGTGCAATTTGGAGCGTTAGCTCGAAAGG<br>GTCTGGATCGGGCCATCACCATCATCACCATTAAtaacaaagctagcataaccccttggggcctctaaacgggtcttgaggggttttttg |
| RMM | taatacgactcactataggggatctagagccccagggtttaaggaggcccacatATGAGTTCGAACGCAATAGGATTAATAGAGACCAAGGGATAC<br>GTCGCAGCCCTAGCCGCCGCTGATGCGATGGTTAAAGCCGCCAATGTTACCATTACAGATAGACAACAAGTAGGTGATGGGTTG<br>GTTGCCGTGATCGTCACAGGTGAGGTTGGGGCAGTGAAAGCAGCTACGGAAGCTGGCGCTGAGACTGCCTCTCAAGTAGGAGA<br>GCTCGTGTCGGTACATGTGATTCCCAGACCGCATTCGGAGCTAGGTGCCCACTTTAGCGTAAGCTCGAAGGGCAGTGGCTCTGGA<br>GCGGACTATAAAGATGACGACGATAAGTAAagggacgcccccaaggagagcaacctATGAGCAGTAACGCCATTGGTCTTATAGAGACC<br>AAGGGTTATGTGGCTGCACTGGCCGCAGCAGATGCTATGGTCAAAGCGGCTAACGTAACGATCACTGACCGACAACAAGTGGG<br>TGATGGTCTTGTGGCAGTCATTGTCACTGGCGAAGTTGGAGCCGTTAAGGCGGCAACAGAAGCAGGCGCGGAGACGGCAAGCC<br>AGGTAGGTGAATTGGTAAGCGTCCATGTCATCCCAAGGCCACATAGCGAACTGGGCGCGCACTTCTCAGTGTCTTCAAAAGGAA<br>GCGGTAGCGGGCATCATCATCACCATCATTAAtaacaaagctagcataaccccttggggcctctaaacgggtcttgaggggttttttg |
